# Lectocyte secrete novel leukolectins *in ovo* for first-line innate immunity defence

**DOI:** 10.1101/2022.07.29.502062

**Authors:** Mirushe H. Miftari, Bernt T. Walther

## Abstract

Atlantic salmon hatching fluid (HF) contains numerous polypeptides. A component unidentified by proteomics, was cloned from tryptic peptides and characterized as lectin-like (LL-) proteins in the tectonin-family. Purified salmon LL-proteins elicited high-titre, LL-specific polyclonal antibodies. This study aims to delineate the cellular and genetic basis of fish embryonic LL-expression. LL-proteins were detected in salmon, cod, rainbow trout and zebrafish HFs. LL-immunoreactive cells were numerous in salmon and rainbow trout embryos, but fewer in zebrafish, cod and halibut. Peridermal salmon LL-positive cells (lectocytes) corresponded to non-eosinophilic cells stained by PAS-reagent. Northern blots revealed two transcripts in salmon and zebrafish embryos, and LL-transcripts were detected specifically in lectocytes. Dual *in situ* hybridization distinguished lectocytes from hatching glands. BAC-library screening yielded salmon Leukolectin’s gene-structure with 4 introns, 5 exons, TATA-box, multiple upstream putative transcription-factor binding-sites, and polyadenylation site. Sequence-analysis indicated zebrafish LL’s conserved nt-sequences and gene-structure, which exhibited mature and truncated LL-transcripts. Zebrafish LL-expression was detected at 6 hpf (yolk syncytium) and 19 hpf (lectocytes and PVF). In dermal mucus, Leukolectins with TECPR-domains may function as pathogen-recognition receptors in first-line innate immunity defence.

**SUMMARY STATEMENT:** At hatching, embryos lose maternal chorions, their first-line innate immuno-protection. Novel leukolectin-genes specifically expressed in non-eosinophilic peridermal cells (lectocytes) help explain how embryos develop innate immuno-competency to survive as larvae.

## INTRODUCTION

Fish development in perivitelline fluid (PVF) is physically safeguarded by the vitelline envelope (or chorion) at stages from zygote until hatching (Kunz, 2004). The chorion-proteins are synthesized in the maternal liver (Hamazaki et al., 1987; Oppen-Berntsen et al., 1992), and transported to ovaries (Hamazaki et al., 1989; Oppen-Berntsen, 1990a; Hyllner et al., 1991) for deposition around differentiating oocytes (Modig et al., 2007). At fertilization, the chorion is polymerized by isopeptide-crosslinks (Oppen-Berntsen et al., 1990b) which essentially abolishes the permeability of macromolecules (Lønning et al., 1984). Molecular passage through the chorion becomes limited to sizes < 4 kDa (Pelka et al., 2017).

Fertilization activates the oocyte (Mei et al., 2009) which causes secretion of maternal (glyco) -proteins deposited in oocytic granules during oogenesis (Ginzburg, 1968; Guraya, 1982; Kunz, 2004; Fuentes and Fernándes, 2010). Due to polymerization of chorion-proteins, such components remain in the PVF and cause its higher osmolarity whereby pliable oocytes are expanded into spherical fish eggs (Lønning et al., 1984; Groot and Alderice, 1985). Thus, the PVF becomes an extra-embryonic compartment with pre-specified macromolecular contents. Both in mechanical and functional respects, the piscine PVF constitutes an embryonic fluid compartment analogous to such compartments in higher vertebrates.

The new appreciation of the functional importance of the PVF has revealed surprising species-diversity concerning PVF-factors and bioactive compounds (De la Paz et al., 2020; Tiralongo et al., 2020), but a generalized PVF-proteome remains to be established. Despite presence in the PVF of immuno-active agents, observations from aquaculture indicate that microbial entry into the PVF after partial digestion of the chorion usually is fatal, suggesting that chorion functions as the essential first-line of embryonic immuno-protection until hatching. If so, the functions of immuno-active PVF-factors are open to reevaluation. Maternal factors contribute to PVF, but developmental changes in PVF during embryogenesis is largely unexplored (Magnadottir et al., 2005). Potential additional roles for PVF-components prior to hatching may relate to functional differentiation of the piscine integument (Elliot, 2011).

Larvae lose both chorion and PVF at hatching (Yamagami, 1988), leaving larvae immunologically challenged by their environment. It is not fully clarified how embryogenesis endows fish larvae with sufficient immuno-protection for life post-hatching. Vertical transfer of maternal defensive molecules to the embryos occurs (Wang and Zhang, 2010; Wang et al., 2012; 2020), but targets for physiological actions of maternal factors are not comprehensively studied. A role in fighting pathogens *in ovo* is possible, but a mode of action preparatory to larval life post-hatching is not inconsistent with assembled data. Particularly for marine fish larvae, very high larval mortality-rates are typical in aquaculture. In contrast, Atlantic salmon larvae normally attain nearly complete (> 95 %) survival. The basis for such discrepant life history outcomes is complex, but nutrition and pathogens are of paramount importance.

Adult fish rely on dermal mucus as their primary first-line of immuno-defence (Ingram, 1980; Esteban, 2012). However, our present understanding of the ontogeny of immuno-competency is incomplete. Immuno-relevant factors may integrate in the dermal glycocalyx of embryos, which post-hatching contributes innate immuno-protection for larvae. The transition from chorionic immuno-protection of embryos to larval immuno-protection by the mucoid glycocalyx remains to be fully delineated.

Specialized cells in fish dermal epithelium secrete a viscous liquid (colloid) with mucins, which produces the mucous layer (Ingram, 1980). The mucus provides a physical barrier against pathogens, which is enhanced by its continuous sloughing-off along with attached pathogens (Rao et al., 2015). A myriad of additional factors such as lysozyme, IgGs, complement, lectins, C-reactive proteins, proteases, protease inhibitors, anti-bacterial polypeptides, alkaline phosphatase, L-amino oxidase, a.o. (Lund and Olafsen, 1998; Wu et al., 1999; Hancock and Scott, 2000; Ellis, 2001; Fast et al., 2002; Birkemo et al., 2003; Matsushita et al., 2004; Roussel and Delmotte, 2004; Bergsson et al., 2005; Luders et al., 2005; Wang and Zhang, 2010; Wang et al., 2012; Arasu et al., 2013; Rakers et al., 2013; Dash et al., 2018) are associated with the dermal glycocalyx, which in sum likely create a bulwark against invading pathogens.

Lectins were discovered in plants and perceived as plant-characteristic proteins before their presence in animals was recognized (Kilpatrick, 2002; Sharon and Lis, 2004). New lectin-like proteins (Huh et al., 1998) with tectonin-structure (Chen et al., 2011) have been demonstrated in fish (e.g., Bildfell et al., 1992; Tateno et al., 2001; Jung et al., 2003; Dong et al., 2004). Their specific immuno-functions are starting to be understood, such as their role in opsonization and phagocytosis (Wang et al., 2016; 2020). The many families of lectins differ vastly in protein-structure, yet all presumably exercise their functions by specific binding to carbohydrates (Sharon and Lis, 2004). Recent studies demonstrated that a group of fish egg lectins (or FELs) specifically recognizes rhamnose (Tateno et al., 2001) and confers binding-affinity for Gram-positive and Gram-negative bacteria. Such interactions do enhance phagocytosis by macrophages (Wang et al., 2020). However, the advent of such processes during embryogenesis awaits clarification.

Modern (salmon) aquaculture entails fish development on an industrial scale, and also makes available a bounty of bio-materials from throughout development. Bio-prospecting hatching fluid (HF) from Atlantic salmon revealed novel proteins, including the endoprotease zonase (Rong, 1997; Walther and Rong, 1998; Miftari et al., 2022) and leukolectins (Miftari, 2009; Walther and Miftari, 2009). Leukolectins (LLs) were discovered during purification of zonase by their affinity-binding to carbohydrate-matrices (Walther and Rong, 1998). LL-proteins are 5-bladed β-propeller proteins (Miftari, 2009; Walther and Miftari, 2009) of the tectonin-family, leading to their proposed role for leukolectins in vertebrate immunity (Miftari, 2009; Miftari and Walther, 2009).

The aims of this work were to establish the occurrence of leukolectins-proteins in various fish species, elucidate the cellular origins of LL-proteins found in hatching fluid, and identify leukolectin’s gene-structure including regulatory elements.

## RESULTS

### Expression and secretion of LL-proteins by peridermal embryonic lectocytes

HF from salmon and cod was analyzed by SDS-PAGE, revealing a complex protein profile after silver-staining (**Fig. 1A)**. Western blot analysis using the diluted specific anti-salmon LL IgG, indicated the presence of one singular molecular moiety (MW around 26 kDa) in both salmon, rainbow trout and cod HF. This finding was supported by two control experiments: The (larger) choriolysins in HF were detected by anti-pike HE antibodies (positive control), while LL was not detected (negative control) in cod without the primary LL-antibody (**Fig. 1A**).

**Fig. 1.**
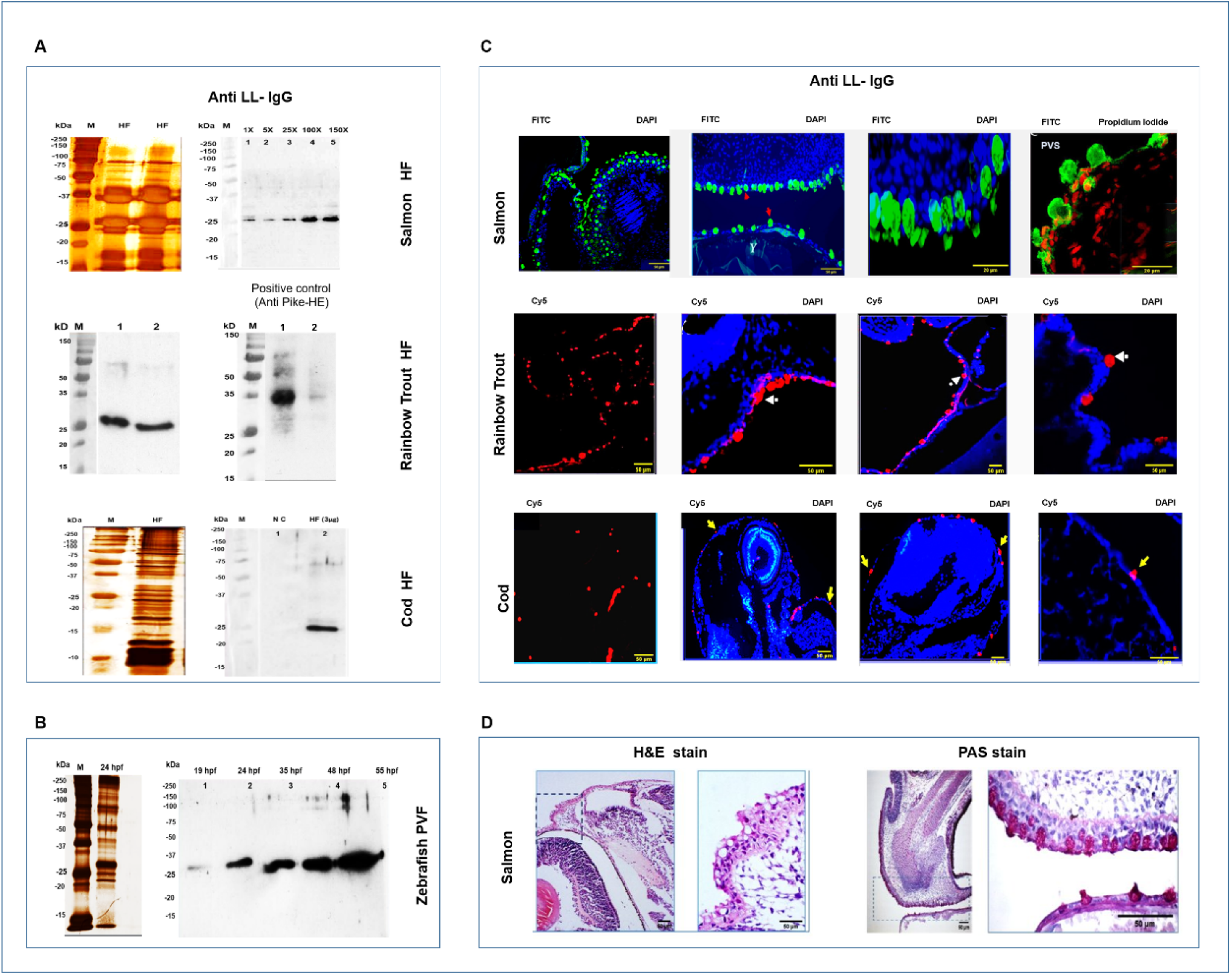
Fish embryonic lectocytes synthesize and secrete LL-proteins. **A**: **SDS-PAGE and Western blots of HF.** <u>Top panels, Left:</u> Two silver-stained salmon HF-samples. <u>Righ</u>t: Western blots of increasing amounts of HF-proteins, showing 26kDa immuno-reactive band. <u>Middle panels, Left:</u> Western blot of two rainbow trout HF-samples. <u>Right:</u> Positive control: Western blot of HF (rainbow trout) immuno-stained with anti-pike HE. Lane 1: high protein amounts show major 35 kDa choriolysin and weak unspecific staining. Lane 2: low protein amounts show 35 kDa band. <u>Bottom panels, Left</u>: Silver-stained cod HF. <u>Right</u>: Western blot of cod HF. Lane 1: negative control (NC). Lane 2, shows single LL-positive band. M (BioRad MW-standard). **B: SDS-PAGE and Western blots of zebrafish PVF.** <u>Left</u>: Silver-stained PVF-sample from single zebrafish embryo (24 hpf). <u>Right</u>: Western blot of PVF from indicated times post-fertilization, showing increasing amounts of 26 kDa LL-positive protein during development. **C**: **Indirect immunofluorescence.** Embryonic paraffin-sections were immune-stained with polyclonal rabbit anti-LL IgGs, detected by fluorochrome conjugated secondary antibody, viewed in appropriate optical filter-set (FITC, or CY5). Salmon sections analyzed by laser-scanning confocal microscopy, and rainbow trout and cod by fluorescent microscopy. <u>Top panel</u> (salmon). First image: Overview of immuno-reactive (green) cells showing numerous peridermal cells. Second image: Arrows (red) point to (green) lectocytes in embryonic and yolk periderm. Note mushroom shapes. Third image: Higher magnification shows vesicular contents in lectocytes (of apparent size differences due to confocal microscopy). Fourth image: Magnified view shows lectocytes with large vesicles projecting into the PVS. <u>Middle</u> <u>panel</u> (rainbow trout), First image: Overview of immunoreactive (red) cells without counterstain. Second and third images: Arrows point to peridermal lectocytes. Fourth image: Magnified view of lectocytes. Arrow points to granular cytoplasm. <u>Bottom panel</u> (cod), First image: Overview, showing fewer immuno-stained cells (without counterstaining). Second and Third images: Arrows point to peridermal lectocytes. Fourth image: Arrow points to lectocyte with typical cytology. Scale bars and counterstain (Dapi, or Propidium iodide) indicated. Y (Yolk), PVS (Perivitelline space). **D: Non-specific staining of salmon embryonic sections.** Left images: Hematoxylin-Eosin staining. Note numerous empty-looking peridermal cells. Magnified boxed area shows non-eosinophilic (empty-looking) cells. <u>Righ</u>t images: PAS-staining reveals numerous positive cells located where non-eosinophilic peridermal cells are seen. Apical domain of PAS-stained cells projects into the PVF, reminiscent of immuno-stained lectocytes (**C**: Top Panel, Right).

We investigated samples collected from zebrafish PVF to eliminate environmental contaminants (**Fig 1B**). Zebrafish PVF presumably does not contain environmental peptides larger than 4 kDa (Pelka et al. 2017). Silver-staining reveals a complex protein-mix in PVF. However, only a single 26 kDa LL-moiety was detected in PVF using anti-LL IgG. This moiety was present as early as 19 hpf, but LL-contents increased during later embryogenesis up to 55 hpf. Apparently, LL-proteins in HF derive from the embryo, and not from environment (**Fig. 1B**). Hence, sections of salmon embryos were investigated by immunohistochemical methods to identify cells synthesizing LL-proteins.

In salmon, indirect immunofluorescence revealed numerous immuno-reactive cells filled with LL-proteins (**Fig. 1C**). These large cells were integral to the periderm (Bouvet, 1976) with long axis oriented vertically towards the PVS. Their apical domain extends beyond the dermal surface as promontories in the PVS. This morphology differs from HG-cells (Schoots et al. 1983a). Such LL-positive cells are seen over the embryo proper, and also over the yolk sac (**Fig. 1C**). Apical domain possesses limited numbers of large (secretory) vesicles (**see Fig. S 1**). The LL-immunoreactive cells were also detected in rainbow trout and in cod embryos, but in lower numbers (**Fig. 1B**). A peridermal location was confirmed, as were their morphological features similar to salmon LL-positive cells (**Fig. 1C**). Even when very few LL-positive cells were present as in halibut, their peridermal location was readily demonstrated (**see Fig. S** 1). In the case for *oikopleura*, LL-positive cells were seen dispersed in the soma (see Fig. S x), and Western blot demonstrated an immuno-reactive (LL-) moiety of the predicted size in *oikopleura* HF (**see Fig. S 2**). Hence, these minority cells may also be lectocytes.

### Non-specific staining of LL-immunoreactive cells

HE-stained salmon embryos possess profuse numbers of empty-looking dermal cells, as these are not well stained by eosin (**Fig. 1D**). These non-eosinophilic cells resemble the LL-positive cells evoked mucus-containing cells (Ingram, 1980), and confirmed by positive control with HE-stained HG-cells (**see Fig. S 3**). Non-eosinophilic cells were further investigated using the PAS-staining method (**Fig. 1D**). The results indicate that PAS-stained cells consistently coincide with the LL-positive cells. Cells described by either staining-method both overreach the embryonic dermal surface. These cells were denoted as *lectocytes.* Lectocytes have not previously been identified in the literature.

### Specific cellular expression of LL-transcripts in embryonic salmon

Cells presenting LL-proteins were verified as also expressing the LL-mRNA. We performed PCR-amplification of LL using LL-specific primers (**Fig. 2A**). Amplicons of 670 bp were generated based on cDNA synthesized from RNA extracted from embryos at mid-embryogenesis (300-370 dd), using β-actin as a positive control (**Fig. 2A**). PCR-products were gel-purified, sequenced and cloned into polylinker site of pBlueScript IIKS (+), as described in Methods. Plasmid DNA was used as a template for *in vitro* transcription to construct LL-specific riboprobes. Anti-sense RNA-probe used SpeI restriction enzyme to generate 5’-overhang (using T3-promoter site), while XhoI was used similarly to generate sense-probe (using T7-promoter site).

**Fig. 2.**
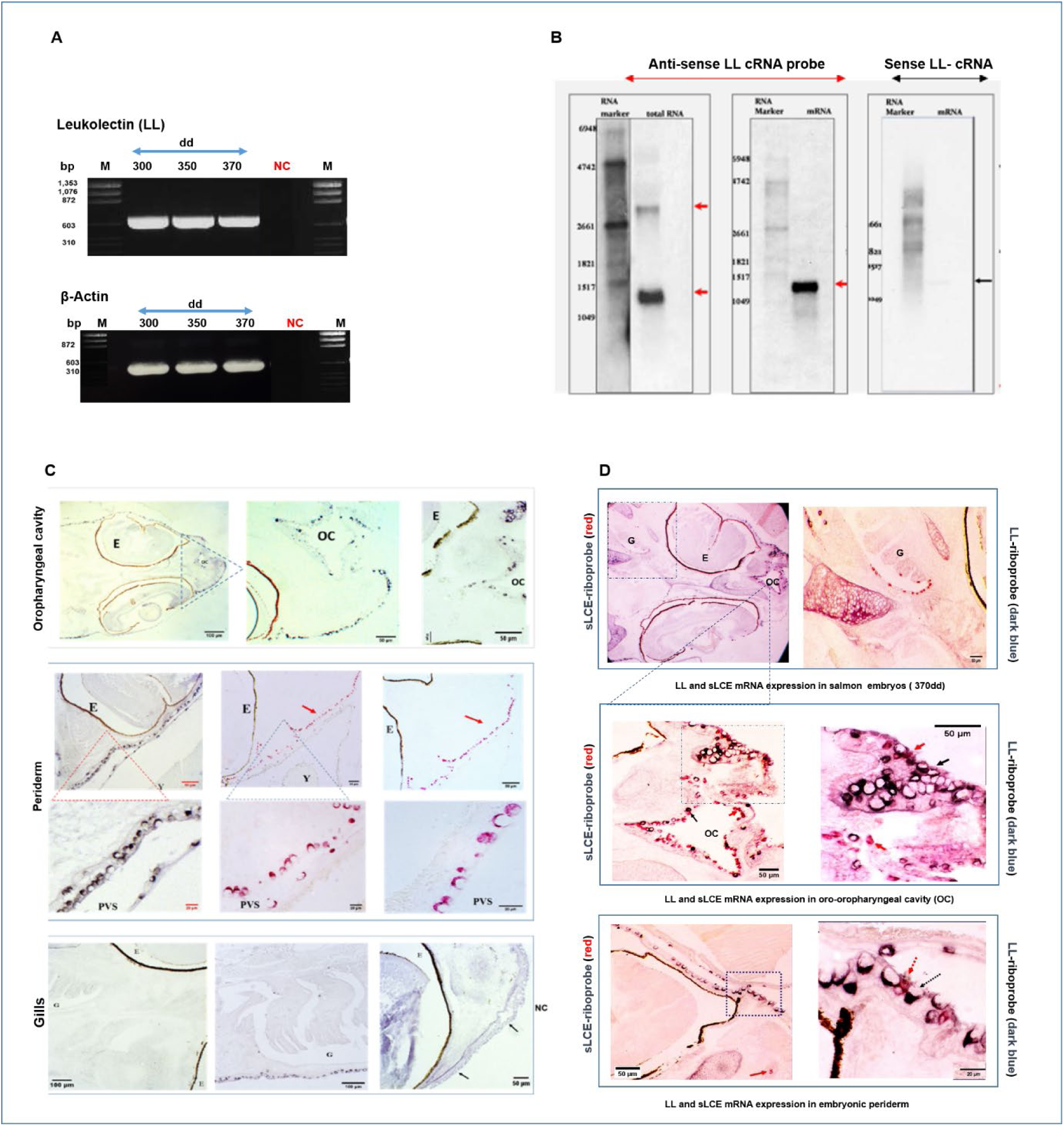
Characteristics and cellular expression of LL-transcripts in salmon embryos. **A: PCR amplification of LL.** <u>Top image</u>: Ethidium bromide-stained agarose gel (1.2 %) showing amplicons (̴ 700 bp), generated with template of cDNA synthesized from total RNA extracted from indicated embryonic stages (in dd). Negative control (NC) shown. <u>Bottom</u> <u>image</u> shows positive control. Amplicons of β-Actin generated using the same cDNA templates. NC (negative control), Marker (M): ϕX174 DNA-HaeIII Digest (NEB). **B: Northern blots analysis.** RNA samples resolved by electrophoresis, were probed with LL-specific DIG-labeled riboprobes (620 bp), see Methods. <u>Lef</u>t image: Northern blot using total RNA from salmon embryos (at 370dd) hybridized with antisense-LL DIG-labeled riboprobe. Red arrows point to the intense minor transcript and weaker larger transcript. <u>Middl</u>e image: Analysis of the poly(A+) RNA fraction coupled to oligo-(dT). Blots hybridized with antisense LL DIG-labelled riboprobe. Red arrow points to single transcript. <u>Righ</u>t image: no hybridization signals detected using sense-LL DIG-labeled riboprobe. Three nitrocellulose membranes shown, with Roche’s DIG-RNA marker I, 0.3-6.9 kb (in left lanes). **C: Single *in situ* hybridization** (*ISH*) **reveals LL-mRNA expression in salmon lectocytes.** *ISH* with DIG-labelled anti-sense LL-riboprobe, detected by anti-DIG antibody carrying alkaline phosphatase (AP). Chromogen used depended on preponderance of endogenous (melanin) pigmentation in tissue-sample, either Fast Red chromogen (insoluble red product), and or NCBI/BIC chromogen (insoluble dark blue product). <u>Top three images</u> show lectocytes in the presumptive oral-pharyngeal cavity (OC). Triangled area reveals singular small and dispersed LL-positive cells within mucosal layer. <u>Middle six images</u>: Top row shows LL-expression (red arrows) in periderm of both embryo proper and yolk sac. Bottom row: Magnified views show large cells (from two triangled areas) with intense LL-expression. Staining restricted to basal cellular portions. Apical domains towards PVS appear empty. A few, smaller cells with uniform cytoplasmic stain seen in periderm. <u>Bottom Left two images</u> of <u>p</u>resumptive gill areas, show a few LL-positive cells. <u>Bottom Right image:</u> negative control (NC) using labelled sense-LL riboprobe (No staining observed). Black arrows point to unstained cells (presumed to be lectocytes). **D: Dual *ISH* shows relative expression of LL and sLCE in salmon embryos.** In formalin-fixed, paraffin-embedded salmon embryonic sections, LL and sLCE transcripts were detected by dual *ISH* using DIG-labeled LL-specific riboprobe (dark blue), and fluorescein-labeled (red) sLCE-riboprobe. <u>Top left image</u>: Cross-section through embryo, with LL-transcripts detected in embryonic periderm, in presumptive OC (triangled area), and in gills (G) in boxed area. <u>Top Right image</u>: Higher magnification from boxed area, with (red) HG-cells in gills, with fewer dark blue LL-cells present. <u>Middle images</u> show magnified views of (triangled) OC-area. <u>Left</u> image: Both gene transcripts are detected, HG-cells (red arrow) being more numerous than LL-cells (black arrow) in mucosal linings. <u>Right</u> <u>image</u> (Magnification of Box in Left image) shows copious lectocytes in periderm (black arrow) with LCE-expressing HG-cells (red arrow) in between, but without co-expression. <u>Bottom images, Left:</u> Periderm with both gene transcripts detected, but with large (dark blue) LL-positive cells (black arrow) predominating, while red (HG-) cells (red arrow) are fewer. <u>Right image:</u> Magnification of Box in Left image. Some dark blue (LL-positive) cells are uniformly stained, while most LL-positive cells are large, and stain intensely only in their basal part. Other cells expressing sLCE transcripts are small and round, and situated between LL-positive cells. <u>Abbreviations:</u> G (Gills), M (Mouth), Y (Yolk), PVS (Perivitelline space), OC (Oro-pharyngeal Cavity), E (Eye, with melanized layers).

Northern blot analysis was applied to define transcripts of LL-genes in salmon, employing a Digoxygenin (DIG)-labeled anti-sense riboprobe (620 nt) specific for LL. The negative control relied on DIG-labeled sense riboprobe (620 nt) (**Fig. 2B**). The positive control applied DIG-labeled anti-sense riboprobe specific for β-actin (data not shown). Samples of total RNA extracted from salmon embryos (370 dd) were resolved by electrophoresis on agarose gel and transferred to nitrocellulose membrane and hybridized with the LL-specific probe (**Fig. 2B**). Two transcripts were detected, including an intensely-staining minor transcript (̴ 1250 nt) and weaker-staining, larger transcript (̴ 2.600 nt).

In order to distinguish between primary and mature transcripts, the analysis was repeated using the poly(A+) RNA fraction isolated from total RNA preparation, by Superparamagnetic Dynabeads®, coupled to oligo-(dT). A single major transcript was revealed (1.250 nt) (**Fig. 2B**), indicating that the larger (primary) transcript (**Fig. 2B**) does not possess a polyA-tail. Control experiments with hybridization utilized the sense-riboprobe, and did not yield a hybridization signal (**Fig. 2B**).

Given LL-mRNA synthesis in the salmon embryo, *in situ* hybridization was employed to paraffin-sections of formalin-fixed embryos. LLmRNA-expressing cells were detected by DIG-labeled LL-riboprobe, with subsequent visualization. This method demonstrated predominant LLmRNA-expression in the periderm, moderate expression in presumptive oral mucosal tissue, and significantly lower expression in epithelium of the gills (**Fig. 2C**). Notably, LLmRNA-staining is restricted to basal portion, with empty apical domains towards PVS. Negative control relied on sense-RNA probe giving no hybridization signal. Some cytological differences between LL-positive cells in oral epithelium (being small and round) compared to large and elongated LL-expressing cells in the periderm (**Fig. 2C**).

Since distribution pattern of LL-positive cells overlaps the known presence of hatching glands in salmon (Miftari et al., 2022), we investigated the distribution pattern of HG-cells and LL-positive cells by use of double *in situ* hybridization (**Fig. 2D**). DIG-labeled LL-specific riboprobe, and fluorescein-labeled sLCE-riboprobe allowed such distinctions in paraffin-sections. In the periderm, LLmRNA-expressing cells are more numerous compared to their presence in the oral region. Also, individual sLCE-positive cells are interspersed between LL-positive cells. This is in contrast to the oropharyngeal epithelium, where HG-cells (expressing sLCE) are more prevalent, and comparatively fewer LL-positive cells are observed. In sum, similar numbers of these two cell-types are found the oral pharyngeal cavity (**Fig. 2D**). In the gills, the LL-numbers are low, while the HG-cells predominate. Even though the two signals were evident in several embryonic tissues, the signals always originated independently from different cells. No co-expression is seen.

### Establishing salmon LL-gene structure

LL-genes were determined by screening Seg.1 High density filter set-007193 from salmon BAC library deposited on CHORI-214, using Digoxigenin-11-dUTP-labelled 620 bp LL-cDNA probe (**Fig. 3A**). The probe is shown (**see Fig. S 4**). Three specific double spots were detected in BAC filter 02D/ Barcode 037214/ Plate 0049/ Range 0096 (**Fig. 3A, left panel**). Also, two specific double spots were detected in BAC filter 03D/ Barcode 037256/ Plate 0097/ Range 01444. (**Fig. 3A, middle panel**). Here, a lower magnification presents the entire filter underneath 4×4 grids. Two specific double spots were identified in BAC filter 06D/ Barcode 037328/ Plate 0341/ Range 0288 (**Fig. 3A, right panel**).

**Fig. 3.**
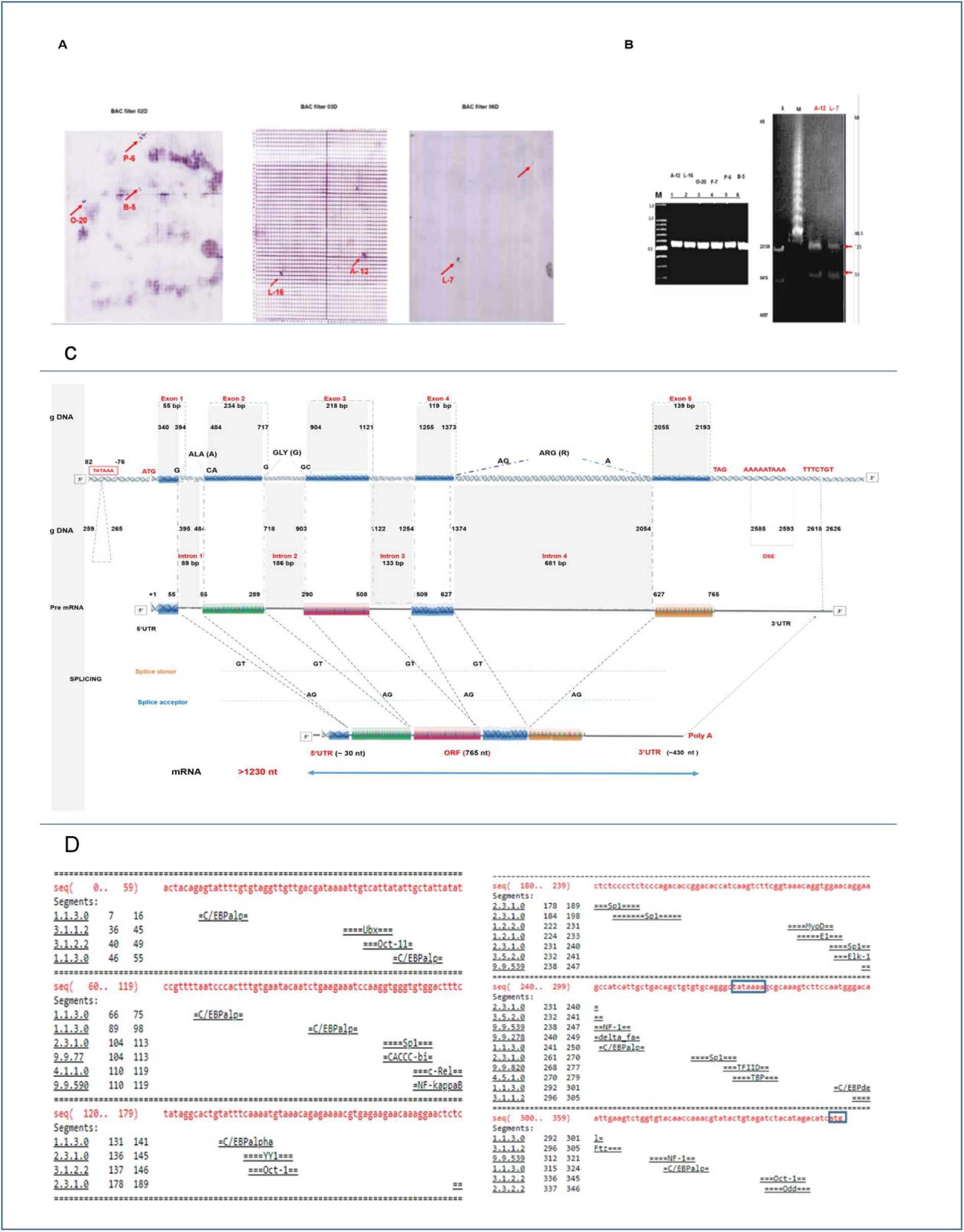
Structure of salmon LL-gene, including upstream and downstream regions. **A: Detection of LL-positive clones in BAC library**. Seg.1 High density filter set-007193 from CHORI-214 salmon BAC library screened using Digoxigenin-11-dUTP-labelled LLcDNA-probe (620 bp). Seven specific duplicate spots (red arrows) detected. On Filter 02D (three spots), on 03D (two spots: L7 & A12, according to BAC-library codes), on 06D (two spots). Middle panel displays entire 4×4 grid filter. **B: Analysis of LL-positive BAC-clones** resolved by gel electrophoresis in Ethidium bromide-stained agarose gels. Left: PCR amplification of LL-gene using purified BAC plasmid DNA as template with LL-specific primers (1.2 % gel). Lanes 1-6 represent six amplicons from six identified BAC-clones, as indicated. M: 100 bp DNA-ladder. Right: Ultra-purified DNA (from clones A12 & L7) after *Not* I -digestion and separated by pulse field gel electrophoresis (1.8 % gel). Red arrows point to two products: 25 kB (insert size) and 13 kB (pTARBAC2.1 plasmid of 13.397 bp). Two markers are: λ-DNA/Hind III-fragments (λ), and NEB’s Low Range PFG-marker (M). **C: Structure of a salmon LL-gene.** Sequence-analysis of one L7-contig (7.080 bp) reveals the complete LL-gene structure. Splicing sites define sizes of 5 exons and 4 introns with split codons for Ala, Gly and Arg. All four splice donors and acceptors are identified as GT and AG, respectively. Positions of start and stop codons, TATA-box, polyA-signal and DSE are shown. An ORF of 765 nt is defined, with 5’-UTR (̴ 100 nt shown) and 3’-UTR of 430 nt. Numbers refer to positions in contig, respectively in cDNA. A primary transcript size between ̴ 2.500-3.000 nt is predicted, with a mature transcript > 1230 nt. **D: An upstream region of the L7-gene** of 359 nt is shown, with TATA-box indicated (Blue Box). Putative TF-binding sites in upstream region were searched by Transfac 4.0 matrices. Multiple binding-sites for numerous TFs are predicted, for a total of some two dozen (overlapping) sites. Abbreviations for the individual TFs are given according to standard nomenclature, and include C/EBPα (7 copies) C/EBPde (1 copy), SP1 (5 copies). UBX (1 copy), NF-Kappa β (1 copy) and NF1 (2 copies).

Seven positive BAC-clones were identified, and six clones were further analysed by PCR-amplification using purified plasmid DNA from each clone as a template, in combination with LL gene-specific primers (**Fig. 3B**). The expected amplicon-size (around 600 bp) were amplified. Lanes 2-7 represent each of six identified LL-positive BAC clones, with marker in lane 1 (**Fig. 3B, left**). The identities of these clones follow from the BAC-library nomenclature, and the various E. *coli*-colonies were cultivated for purification of plasmid DNA. From two BAC clones (L-7 and A-12) purified plasmid DNA was digested with *Not* I restriction enzyme, and separated by pulse field gel electrophoresis with Ethidium bromide-stained gels. The data show (**Fig. 3B**) two bands, one with MW of 13 kbp (corresponding to the pTAR BAC2.1 of size 13.397 bp), and a second of MW of 25 kbp. Shown (**Fig. 3B, right**) are the inserts derived from digested BAC clones A12/142 and L7/257.

Shotgun sequencing by MRW (Berlin, Germany) of the 25 kbp DNA-inserts from these two BAC clones (denoted as L-7 and A-12) resulted in a total of 80 reads, which were assembled into 33 contigs. Here, the LL-gene structure is reported from the sequence analysis of one contig which derived from L-7 BAC-clone (**see Fig. S 3B**), and summarized results are presented (**Fig. 3C**). The coding region for LL encompasses five exons (of respective lengths 55 bp; 234 bp; 218 bp; 119 bp, and 139 bp), and four introns with respective lengths (89; 186; 133; 681 bp) as outlined (**Fig. 3C**). There are three identified split codons coding for Ala (between Exon1 and Exon2), Gly (between Exon2 and Exon3), and Arg (between Exon4 and Exon5). Organization of LL-gene from this contig, includes an upstream region (up till −360 bp), TATA-box (in position −72 – 80, relative to the start codon ATG. The 3’-UTR follows stop codon TAG, and spans around 440 bp, with a DSE -signal at position 2585-2593, and a polyA-site at 2618-2626. The primary LL-transcript is at least 2700 nt, while the mature transcript is larger than 1230 nt (**Fig. 3C**). The splicing donor and acceptors are identified in all four positions.

The upstream region of the LL-gene (**Fig. 3D**) displays multiple putative transcription factor (TF-) binding sites. The various TF-binding sites for c/EBPα, for NF-κB and for SP1 are indicated. Further binding-sites were also predicted for OCTα, YYx, and MynD. Many such binding sites are found in multiple copies. Since binding sites of important regulatory transcription-factors occur in the upstream region of salmon LL-gene, a BLAST-search for other genes involved in innate immunity in the LL-positive contig was performed. The MHC II gene with innate immunity function (**Harstad et al., 2008**) and IgH (Seq. ID GU129139.1) were found to also be present on LL-contig (**Fig. S 5**). Both the MHC-II and the IgH genes are encoded in the 3’-direction on the LL-contig (**Fig. S 5**). The LLgene-sequences were deposited in GeneBanks (2009). Salmon leukolectin sequences were deposited (GeneBank accession numbers FN296284.1). In the updated salmon genome-version Ssal_v3.1, the LL gene was located on chromosome ssa 17, while not present on the earlier assembly (Lien et al. 2016).

### Structure of Zebrafish Leukolectins

Salmon is a species with limited history for fundamental scientific investigations, while zebrafish is an acknowledged model species (Nüsslein-Volhard and Dahm, 2002). Also, whole mount RNA *in situ* hybridisation in zebrafish allows surveying endogenous gene expression during embryogenesis. Therefore, zebrafish leukolectins were investigated. Initially, partial coding sequences of zebrafish LL were amplified using salmon LL-genespecific primers (**Fig. 4A**).

**Fig. 4.**
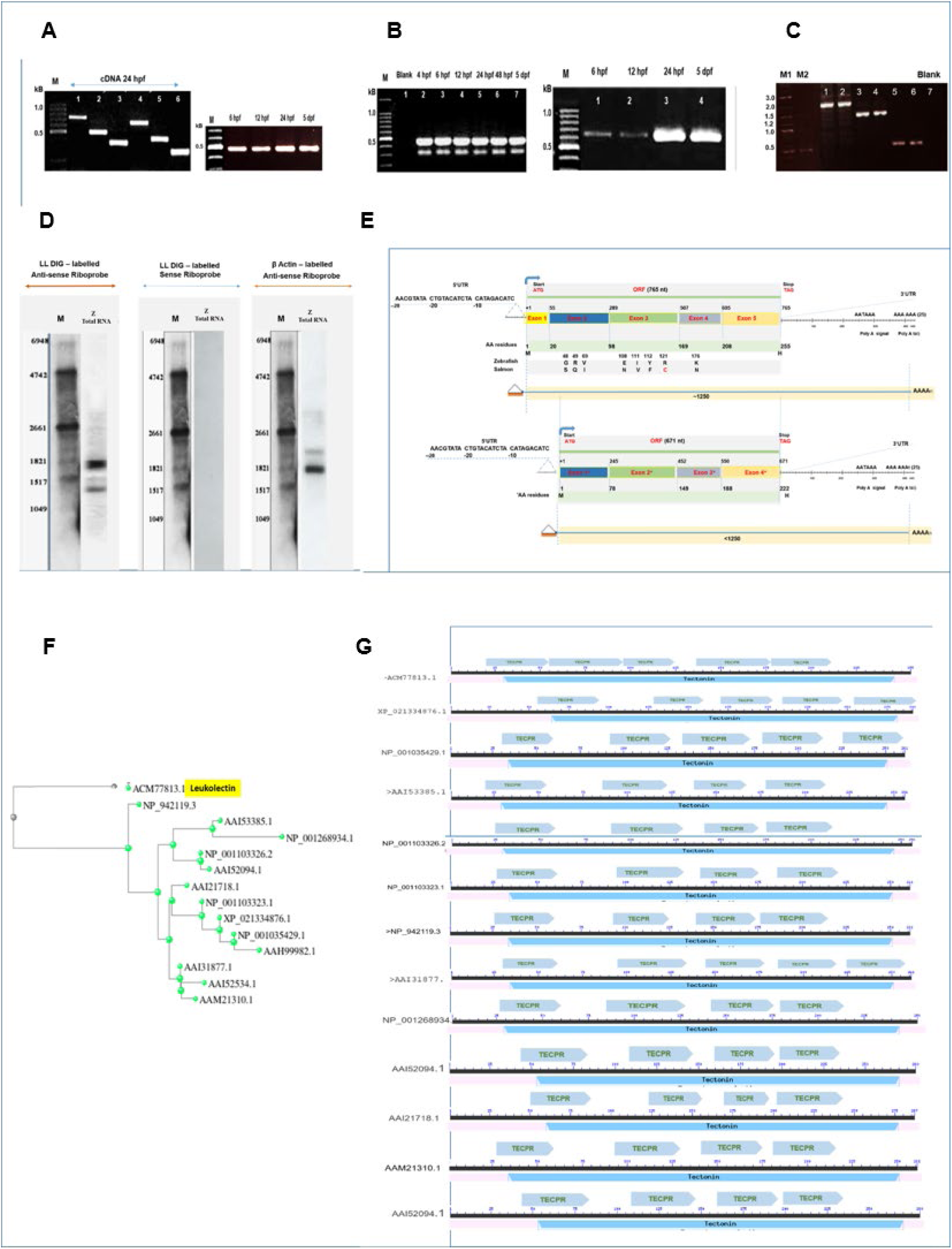
Structure of Zebrafish Leukolectins (zLL) **A: PCR Amplification.** Amplicons were generated using cDNA synthesized from total zebrafish RNA at 24 hpf, using six different combinations of (salmon) LLgene-specific primer pairs. <u>Left image</u>: Amplicons resolved in Ethidium bromide-stained agarose gels (1.2%). Lanes: 1 (720 bp), 2 (510 bp), 3 (350 bp), 4 (650 bp), 5 (450 bp), 6 (280 bp). <u>Right image</u>: Positive control with β-Actin gene, using cDNA from various embryonic stages. **B: RACE-PCR**. <u>Left image</u>: Two amplicons generated by 5’-RACE PCR from the indicated stages, respectively around 500 and 350 bp. The smaller amplicon is present in significantly lower amounts. <u>Right image</u>: a single amplicon of around 650 bp was generated using cDNA from different developmental stages by 3’-RACE PCR. **C: PCR Amplification.** Amplicons generated using DNA extracted from 24 hpf embryos. Forward primers were designed based on salmon (5’-UTR) genomic LL-sequence, and Reverse primers positioned < 1000 bp apart to allow direct sequencing of the gene-portions. Amplicons analysed in duplicates. Lane 1-2: about 2.000 bp. Lane 3-4: > 1.500 bp. Lane 5-6: around 700 bp, Lane 7: Blank. M1 (NEB’s 2-Log DNA Ladder), M2 (TRAN’s 100 bp DNA Ladder). **D: Northern blot analysis**. Total zebrafish RNA (zRNA) was extracted from embryos at 24 hpf, and probed with anti-sense zLL DIG-labelled riboprobe (620n t). <u>Left Image</u>: Two transcripts detected of 1.300 nt (at arrow) and 1.800 nt. (Truncated transcript not detected). <u>Middle image</u>: No transcript detectable using sense-zLL DIG-labelled probe. <u>Left image:</u> Positive control using anti-sense β-Actin DIG-labelled riboprobe. **E: Summary of zebrafish LL transcripts.** <u>Top panel</u>: Mature full-length LL-transcript with ORF (765 nt), 5’-UTR of 28 nt and 3’-UTR of around 420 nt (total size ca.1.235 nt) are shown. Five exons and their sizes are indicated. A comparison with salmon LL shows positions of only eight differing AA-residues (two major changes). 5’-UTR is shown in full, and positions of polyA-modification sites are indicated in 3’-UTR. <u>Bottom panel:</u> Smaller (truncated) transcript (from Fig. 4B) of 671 nt, starting at Met(M) in position 32 in mature transcript, are shown. The approximate size the truncated form is around 1160 nt. Its constituent four exons correspond to Exons2-5 in the full-length LL-transcript. 5’- and 3’- ends are identical compared to full-length transcript. **F: Phylogenetic tree for zebrafish lectins.** Constructed by the neighbour-joining method, with maximal sequence difference of 0.6. Query sequence was zLL (ACM77813.1). Sequence IDs of thirteen Fish Egg Lectins (FELs) in the zebrafish database are indicated. LL forms a separate protein branch, and while twelve other FELs branch from the NP_942119.3 entity (Pasquier et al., 2016). **G: Prediction of putative TECPR-domains in zLL and other zebrafish FELs.** zLL and three other zFELs (XP_021334876.1, NP001035429.1 and AA131877.1) are predicted with five TECPR-domains by SMART database. Note that sequence ID (NP_942119.3) is predicted to possess four TECPR, along with eight other zFELs, despite its distinct branching in the phylogenetic tree (Fig. 4F).

Six amplicons were generated by choice of primer-combinations. Their sizes appeared as predicted, provided zebrafish LL roughly correspond to salmon LL. Positive control relied on β-actin amplification. Amplicons were gel-purified and sequenced, establishing partial LLcDNA coding -sequences. Zebrafish specific primers were designed to generate the full-length cDNA sequence. 5’-RACE-PCR gave amplicons of two sizes (of ca. 500 and 350 kbp) generated using cDNA synthesized from RNA extracted from developmental stages 4 hpf to 5 dpf. The larger amplicon of 500 bp was gel purified and sequenced by direct sequencing. The data reveal the entire 5’-RACE oligoRNA sequence followed by 28 nt representing 5’-UTR before the ATG start codon (first M-residue) continuation with Exon1-sequence (ending at reverse primer). The minor amplicon of 350 bp was also gel purified and sequenced, revealing the same upstream 28 nt, but continuing with nucleotides in the Exon2. This sequence also starts with methionine, which is the first M-residue in Exon2. Apart from the initial 5’-portion, the sequences of mature and truncated mRNAs were identical.

In contrast, 3’-RACE-PCR gave one single amplicon using RNA from 6 hpf to 5 dpf (**Fig. 4B**), and subjected to direct sequencing. The 3’-UTR appears very similar to salmon 3’-UTR, except singular nucleotide-changes. Concerning the zebrafish 5’-UTR, the 28 nt identified were also identical nt in salmon 5’-UTR. Exhaustive sequencing gave the entire nt-sequence of zebrafish LL-mRNA. Analysis of full-length cDNA-sequences reveals a very high overall similarity to salmon LL. A comparison of putative AA-sequences of salmon and zebrafish LL show highly conserved structures, differing at only 8 residues (**Fig. 4E**). The differences are radical at 3 positions only, but one of these sites include substitution of one Cys (in Exon3) with an Arg, which implies that one of the 3 SS-bonds in salmon LL-protein is missing in zebrafish. Full-length LL-cDNA sequences were deposited (Gene accession FJ643620.1), and part of missing cDNA in earlier zebrafish assemblies (Howe et al., 2013).

These results were supported by Northern blot analysis of zebrafish total RNA (**Fig. 4D**). Two transcripts are seen. The major transcript has a size compatible with a presumptive primary transcript, although smaller than the primary transcript seen in salmon. The minor zebrafish transcript (at arrow, **Fig. 4D, left image**) is inferred to be a mature LL-mRNA. Positive control (Right image, using β-actin sense riboprobe) and negative control (Middle image, using sense riboprobe) are presented.

The zebrafish LL-gene structure was established as follows. Genomic DNA (extracted at 24 hpf) was used as a template in combination with one LL-specific forward primer (designed from salmon 5’-upstream gene region) combined with 3 different reverse primers for PCR-amplification. Primer sequences and positions are shown (**see Fig. S 5**). Agarose gels with the amplicons (applied in duplicates) are presented (**Fig. 4C**). Amplicons were gel-purified and sequenced. In sum, the LL-gene structure with 5 exons, 4 introns, and their individual sizes and splicing were identical to salmon (**see Fig. S 6 A**).

BLASTp network server at GenBank database searches revealed a 43-47 % sequence similarity of putative LL-proteins compared to several other deposited zebrafish egg lectins (**see Fig. S 6B**). This allows constructing an evolutionary LL-protein tree, where LL appears as a separate branch, originating separately from the other zebrafish lectins (**Fig. 4F**).

All these zebrafish lectins are predicted as belonging to the tectonin protein family. The domain-structure of putative LL-proteins was investigated using SMART database (**Fig. 4G**). Most of the zebrafish lectin appear with 4 TECPR-domains, while zebrafish LL and three other fish egg lectins (XP_021334876.1 and NP_001103323.1 and AA131877.1) possess 5 TECPR-domains. These lectins possess respectively 45.8 %, 46.7 % and 46.7 % similarity to leukolectins, compared to a 96.9 % similarity of putative salmon and zebrafish LL-proteins.

Additionally, multiple sequence alignment of putative zebrafish lectins reveals conservation of Cys-residue positions. Available sequences allow comparisons to 13 other egg lectin proteins, showing 4 Cys-residues occur in invariant positions, while zebrafish LL alone is missing a 5th Cys-residue present in other egg lectins (**see Fig. S 6B**).

### Cellular expression of LL-transcripts-in zebrafish embryos

Gene expression of LL in zebrafish was investigated using whole mount RNA *in situ* hybridization using a single probe. At early developmental stages (11-12 hpf) LL-transcripts are confined in one cellular type, which are more numerous above the yolk. In sections of these embryos, transcripts are predominately found in the yolk syncytial layer (**Fig. 5A**).

**FIG. 5.**
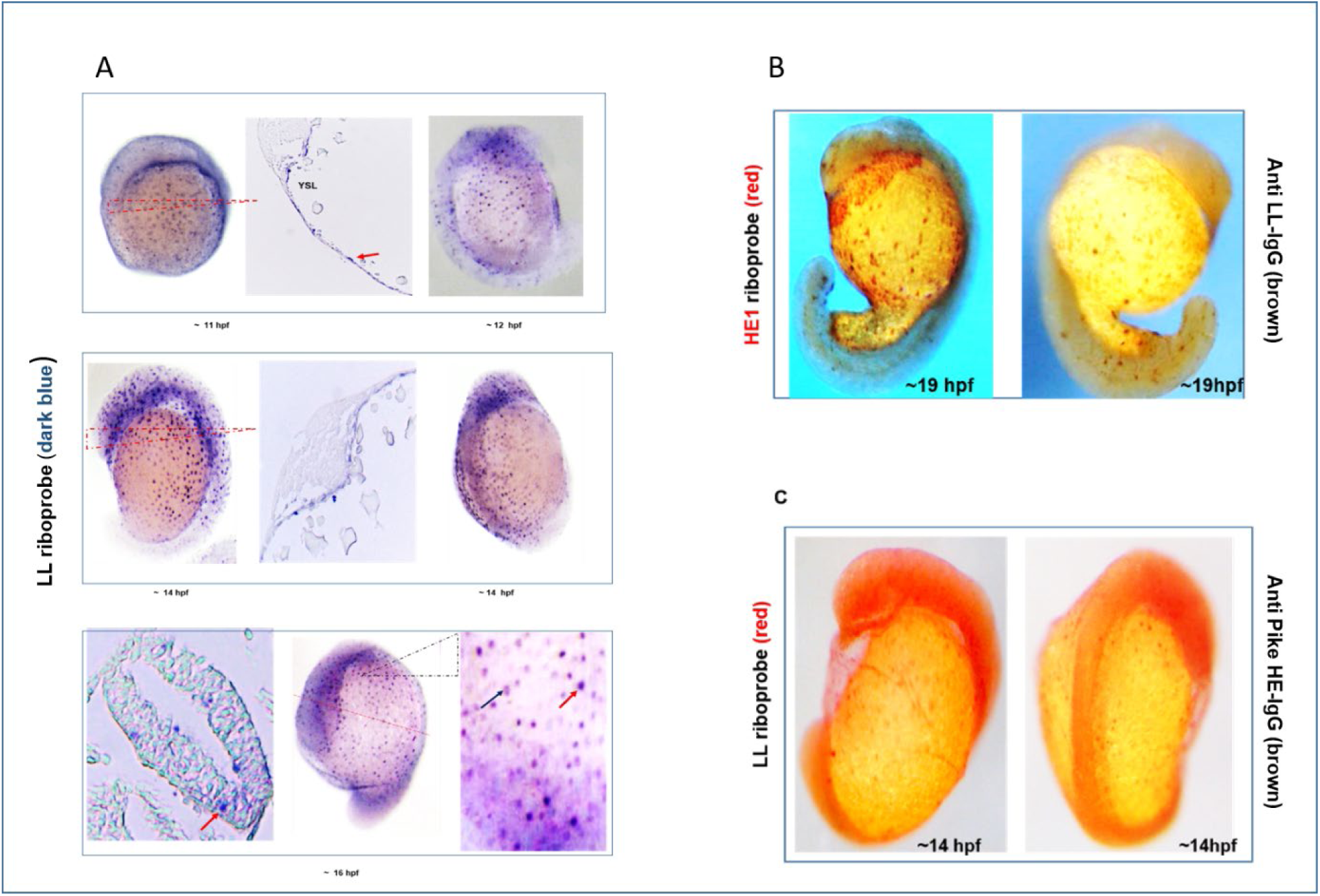
Zebrafish LL and choriolysin are expressed in different cells. **A: Single label whole mount RNA** *ISH* **of zebrafish embryos using anti-sense zLL DIG-labelled riboprobe.** <u>Top panel</u>: Left and Right montage images show early embryos (at 11 & 12 hpf) with (dark-blue) LL-transcripts detected over embryo and yolk sac. Middle image shows cross-section from the triangled area of Left image. LL-transcripts in yolk syncytial layer (YSL) are noted (red arrow), which are mostly absent in the embryo proper. <u>Middle</u> <u>panels</u>: Later embryos (14 hpf) show LL-transcripts in cells found in lateral (Left image) and dorsal (Right image) aspect of embryos. Middle image shows cross-section from triangled area in Left image, revealing LL-transcripts in YSL in distinct cells. <u>Bottom panels:</u> Whole mount of 16 hpf embryo shown. Middle image shows wide distribution of LL-transcripts in embryo proper and over yolk sac. Left image shows cross section (at indicated position in Middle image), with LL-positive cells within the neural tube (red arrow). Right image shows magnified view of triangled area (in Middle image) with intensely stained compact cells (red arrow) and weakly stained cells (black arrow). **B: Whole mount RNA** *ISH* **of zebrafish embryos at 19 hpf.** Anti-sense zebrafish HE1-riboprobe (red) was employed in combination with immunohistochemistry using anti-LL IgG-antibodies (DAB brown). Left image shows HE1-positive (red) confined to the known HG-location (black arrow). In contrast, LL-protein (brown DAB) is expressed in single dispersed cells throughout embryonic tissues. Right image: LL-positive cells display variant staining intensities. Some strongly stained (large and rounded) cells are found in the tail area (black arrow). **C: Whole mount RNA** *ISH* **of 14 hpf zebrafish embryos.** Anti-sense LL-riboprobe (red) used in combination with immunohistochemistry employing anti-Pike HE-antibodies (DAB brown). At 14 hpf, LL-transcripts are found in individual dispersed cells in the embryonic tail and back side, where HG-cells are known to be absent (black arrows). Embryonic HG-cells are confined to pre-chordal plate, and are stained brown DAB-colour.

Already at 14 hpf, more LL-positive cells are found in the embryo proper. At this stage, cellular appearances are diverging. Some cells are numerous and intensively stained, and relatively large in whole mount images. The other cell type is round and seemingly smaller, with weaker staining. Sections of embryos reveal no clear LL-transcripts in the embryo proper at 14 hpf. However, at 16 hpf, transcripts are detected within the embryonic soma. Elsewhere in the embryo, transcription is observed in the above-mentioned two cellular types.

### Cellular expression of LL-proteins and choriolysin-mRNA

In order to substantiate the conclusion concerning distinct identities of lectocytes and HG-cells in early embryos, whole mount RNA *in situ* hybridization using HE1-riboprobe was employed in combination with immunohistochemistry, using anti-salmon LL-IgG (**Fig. 5B**). Again, the data show the distinct identities of (brown LL-proteins in) lectocytes and of (red) choriolysin-expressing HG-cells. Anatomical HG-localization in zebrafish at 19 hpf is well known (Inohaya et al., 1997), and is shown here as a reddish cell group in pre-chordal position over the yolk sac. Focusing on the zebrafish tail at this stage, HGs are known to be absent. Hence, the brown cells in the tail area reflect only LL-positive cells. Thus, at least in the tail-area LL-cells are clearly distinct from HG-cells. The same distinction is possible but less clearcut in other parts of the embryo, where the two cell types occur together, a distinction which is further exemplified (**see Fig. S 7**) by additional data on lectocytes and HG-cells.

### Cellular expression of LL-transcripts and choriolysin-proteins

In order to clearly distinguish between lectocytes and HG-cells in early embryos, whole mount zebrafish embryos were investigated by combined immunohistochemistry (using anti-pike HE) with *in situ* LL-hybridization. Since the embryonic location of HG-cells is well documented in zebrafish, the data (**Fig. 5C**) show that LL-transcripts are seen in places where HG-cells are absent at 14 hpf, and *vice versus*. In sum, not only the cytology but also the anatomical positions of lectocytes and HG-cells are identified in zebrafish embryos.

## Discussion

Lectocytes are embryonic fish peridermal cells which express novel leukolectins-proteins with five TECPR-domains. LL-encoding gene-sequences are highly conserved, with a phylogenetic tree distinct from other fish egg lectins. Salmon LL-transcription appears regulated by multiple pivotal transcription factors. From mid-embryogenesis, lectocytes secrete LL-proteins into the PVF. Leukolectins integrated within dermal glycocalyx may enhance functionality of mucus which will replace the (maternal) chorion as larval first line of immuno-defence after hatching.

Leukolectin-synthesis in embryonic peridermal epithelial lectocytes was shown using specific polyclonal IgGs (Fig. 1C), and confirmed by *in situ* hybridization (Figs. 2, 5) using LL-specific riboprobes. Surprisingly, many peridermal PAS-stained mucous cells in salmon (Fig. 1D) synthesize LL-proteins (Fig. 1C). The differentiated lectocytes are not recognized by routine histological methods, which show embryos to be replete with non-eosinophilic (empty-looking) cells (Fig. 1D). These cells are unstained by eosin (Fig. 1D), despite such cells (lectocytes) being filled with LL-proteins (Fig. 1C). Eosin-staining of proteins may be hindered if pI of proteins is near neutrality (e.g., LL) or if modified by prosthetic carbohydrate-groups. Hence, conclusions from negative eosin-staining in histochemical studies warrant re-examination.

In the piscine dermis, differentiated lectocytes and HG-cells are intermingled, but not co-expressing their respective marker-transcripts (Figs. 2, 5). LL-positive peridermal cells contain ample secretory vesicles (**see Fig. S 1**) prior to the appearance of cyto-differentiated HG-cells (Inohaya et al. 1997). Both HG-cells and lectocytes perform exocrine secretion into the PVF (Fig. 1A, B), and both possess cytology with apical secretory vesicles (Fig. 1C; **See Fig. S 1**), and LL-mRNA assembled in basal RER-compartment (Fig. 2). However, control of secretion by these two cell-types is fundamentally different. HG-cells do not secrete until signals from the environment cause hatching (Schoots et al., 1983b; DiMichele and Powers, 1984; Yamagami, 1988; Oppen-Berntsen et al., 1990c, Helvik and Walther, 1992; 1993a; 1993b), while lectocytes start secreting from mid-embryonic life (Fig. 1). These observations emphasize the distinct categories for factors functioning in early embryogenesis: maternal proteins (including lectins) deposited in oocytes, maternal factors with transcription (included for lectins) needed in embryos before the mid-blastula transition (Kane and Kimmel, 1993), and embryonic genes transcribed during embryonic cyto-differentiation. Both Leukolectin and choriolysin-genes belong in the latter category (Miftari et al. 2022), but LL-genes are expressed earlier than choriolysin-genes (Fig. 4B).

LL-transcripts are found in salmon in mid-embryogenesis (Fig. 2), when both primary and mature transcripts are seen. Transcripts are present in numerous dermal epithelial cells over most of the soma and yolk sac, while lower expression is evident in the mouth and gills. This contrasts with HG-cells (Yokoya and Ebina, 1976; Miftari et al., 2022). Some lectocytes in the dermis and oro-pharyngeal epithelium exhibit variant cytology, for unknown reasons.

Mucous cells (Elliot, 2001) are believed to differentiate within the epithelial dermis. The origins of HG-cells are defined (Inohaya et al., 1997), while stem-cells for lectocytes are not yet defined (Fig. 5A). Mucous cells function in first-line innate immunity defence (Ingram, 1980; Subramanian et al., 2007; Dash et al., 2018), implying that lectocytes (Fig. 1) also participate in this regard. LL-contents may potentiate innate immunity functions of lectocytes. Assuming that lectocytes remain a feature of dermal epithelium *throughout* life, LL-presence in dermal mucus of grown fish could influence responses to pathogens, and perhaps forecast the outcome of pathological challenges. Since many (diverse) lectins are reported in fish (e.g., Tsutsui et al., 2003; Rajan et al., 2011; Cordero et al., 2015), establishing their possible expression in lectocytes, or alternatively, in other distinct mucous cells (Elliot, 2011), is of interest. Conceivably, adaption of fishes to different biotopes may have relied on evolving biotope-relevant innate lectins to achieve immuno-protection.

The data (Fig. 3) show that an LL-gene possesses a distinct, but conventional structure, with features including TATA-box, splicing sites (without alternative splicing detected), exons, introns and signals for a poly-A tail. One distinguishing feature of the salmon LL-gene is its upstream region (Fig. 3D), with multiple (putative) binding sites for several transcription factors. Sp 1-sites suggest a general relevance of LL for processes during growth, development and differentiation (Beishline and Azizkhan-Clifford, 2015). A specific relevance for myelogenesis is indicated by binding-sites for C/EBPα (Friedman, 2007), and for immuno-defence by NFκB-site (Dorrington and Fraser, 2019). Despite our finding of non-expression of (some) other immunity-related markers in salmon lectocytes, these findings offer clues as to identities of non-lectocytic cells expressing leukolectins. LL-transcripts appear in undifferentiated cells fated to become lectocytes, but are also seen in cells within the neural tube (Fig. 5A) with no clear connection to peridermal skin. Also, a few other cells with LL-transcripts possess unusual cytology, which suggests that LL may function is non-dermal cells (Miftari and Walther, 2009; Miftari and Walther, Manuscript under revision).

Salmon and zebrafish LL-genes (Fig. 4) displayed similar features. The zebrafish primary transcript seemed slightly shorter than its salmon counterpart, while mature transcripts exhibited a similar size. Also, a truncated zebrafish LL-transcript was detected without a putative signal-peptide domain. LL-genes in these two fishes show high sequence conservation, with predicted 91 % homology. Zebrafish Fish Egg Lectin (zFEL; Wang et al., 2016) and LL are both predicted to belong to the tectonin-family, with five (or four) TECPR-domains. However, the two lectins showed a sequence identify of only 48 %, placing LL- and zFEL-proteins as separate branches on a phylogenetic protein tree (Fig. 4D).

TECPR-domains (Chen et al., 2011; Qiao et al., 2022) have recently been documented as relevant for lectin-recognition of carbohydrate-moieties and modified (methylated) carbohydrates-moieties (Wohlschlager et al., 2014). Factors controlling immunological first-line defence are still incompletely understood, but recent findings (Stowell et al., 2014) point to such recognition patterns as vital defence mechanisms in multicellular organisms against predators, parasites and pathogens (Cummings, 2014). Lectins recognizing specific sugars, command paramount importance in innate immunity defence (Sommer et al., 2018), and specific roles of individual TECPR-domains in transmembrane signalling during phagocytosis, autophagosome maturation and autophagosome fusion are recognized (Fraiberg et al., 2021). Among tectonin-domains, TECPR3 was predicted to bind bacterial LPS, while TECPRs 2&3 were predicted with galactose-affinities (Low et al. 2010). Also, mutations in TECPR2 domain were found to entail consequences for autophagy (Oz-Levi et al., 2012).

The ‘*Lectin recognizing Glycan’* concept of innate immunity is in line with how structural protein-domains for ‘*Self: Non-Self’* recognition are specified (Sommer et al., 2018). Their evaluation of lectins in higher vertebrates concluded that only leukolectins possess the carbohydrate binding-sites in domains 2 (&3) needed for such interactions. If these conclusions are sustained by further studies, leukolectins constitute crucial defence lectins in innate immunity. Thus, Leukolectin-actions may not be limited to the needs of larval fish for immuno-defence during early development.

## MATERIALS and METHODS

### Biological Materials

Biological samples from the following fish species Atlantic salmon (*Salmo salar* L.), Cod (*Gadus morhua*) and Halibut (*Hippoglossus hippoglossus*) were obtained from the Marine Institute, Bergen, and from AS Bolaks, while Rainbow trout (*Oncorhyncus mykiss*) samples were from Tomre Fishfarm (Fusa). Zebrafish (*Danio rerio* zebrafish) and *Oikopleura dioica* samples were obtained from the SARS Centre, Bergen.

Hatching Fluids (HF) from all fishes were obtained as described (Rong, 1997; Miftari et al., 2022), where HF is collected as soon as at least 50 % of eggs have hatched. Zebrafish perivitelline fluid (PVF) was collected from single embryos placed in 20 µl destilled water during dechorionation. Samples were collected and kept frozen for further analysis.

### Primary antibodies

Polyclonal rabbit anti-leukolectin antibodies were prepared using sequence-grade pure salmon Leukolectin protein (sLL) as immunogen (Rong, 1997). Chinchilla rabbit (4 kg) was immunized at Vivarium, Univ. of Bergen following the immunization protocol (Harlow and Lane, 1988) for six weeks. Initial injection of immunogen included 80 μg of purified protein sample emulsified with 500 μl of Freund’s complete adjuvant. Next, two subsequent injections were performed using 40 μg and 25 μg of purified protein sample mixed with 500 μl of incomplete Freund’s adjuvant according to the manufacturer’s instructions. The immunogen was injected into multiple subcutaneous sites of the rabbit. Total blood volume (150 ml) was collected eight days after the final booster. The serum was prepared after centrifugation at 3,000 rpm (Sorvall SS-34 rotor) for 15 min. Aliquots of antiserum were stored at −80 °C for long storage. Primary antibody against choriolysin (Polyclonal rabbit anti-pike HE antibody) was gift from prof. J. M: Denucé, Univ. of Nijmegen, Holland. For later work, sequence-specific LL-antibodies were designed, which allowed verification of the results from use of polyclonal LL-antibodies.

### Methods

#### IgG fraction purification from the antisera

Primary antibodies were affinity-purified using HiTrap 1-ml Protein G columns (Amersham Pharmacia Biotech) with flow rate 1 ml/min. Initially, the antiserum sample was filtrated through a 45 µm filter unit and applied to 1ml Protein G columns. The column was equilibrated using 5-10 column volumes of binding buffer (20 mM Na-phosphate, pH 7.0) followed by several washes with the same buffer until no protein detects by UV absorbance at 280 nm. Next, IgG fractions were eluted with 5 ml of elution buffer (0.1 M glycine HCl, pH 2.7) and divided into ten tubes containing 25 µl of neutralization buffer (1 M Tris HCl, pH 9.0). Protein concentration was determined spectrophotometrically while the purity was estimated by analyzing the aliquots of purified IgG fractions in 12 % SDS-PAGE with subsequent silver staining. In addition, antibody reactivity was evaluated by Western blot analysis using different dilutions of primary antibody with subsequent detection using goat anti-rabbit IgG, horseradish peroxidase-conjugated secondary antibody. After purification, IgG fractions were passed over though a PD-10 column (Amersham Pharmacia Biotech)) for buffer exchange to 0.1 M Na-phosphate (pH 7.4), 0.15 M NaCl, and 0.1 %, Na Azide. The aliquots are stored at 4 °C or –80 °C for long-term storage. To perform the zebrafish experiments, polyclonal antibodies were pre-adsorbed by incubation in zebrafish acetone powder before the IgG purification procedure (Nüsslein-Vollhard and Dahm, 2002).

#### SDS-PAGE and Western blot analysis

Samples were resolved in 12 or 15 % SDS PAGE (Laemli, 1970) and silver stained in accordance with the protocol (Heukeshoven and Dernick, 1985). Electrophoresed protein samples were transferred to nitrocellulose blotting membranes (Hybond, Amersham Pharmacia Protram 0.45 μm NC) in presence of blotting buffer, under constant voltage (Towbin et al., 1979). Afterwards, the nitrocellulose membranes were incubated in blocking buffer with 3 % gelatin (Sigma-Aldrich) in TTBS buffer (Tris-buffered saline, pH7.5, containing 0.05 % Tween-20) for 30–45 min with agitation. Membranes were incubated in the solution of diluted primary antibodies at 4 °C, overnight. Rabbit polyclonal anti-LL IgG were applied diluted 1: 5.000 in a blocking solution while, rabbit polyclonal anti-pike-HE antiserum was applied diluted 1: 2.000 in the same buffer. Subsequently, the membranes were washed with gentle shaking for 2x 15 min in TTBS before incubating with goat anti-rabbit IgG, HRP-conjugated secondary antibody (Dako), diluted 1: 2.000 in blocking buffer, at RT, for 1 h with agitation. The membranes were washed twice in 1x TTBS (15 min) and twice in 1x TBS (5 min). Detection in immunoblots relied on an enhancer substrate (Super Signal West Pico Chemiluminescent Substrate (ThermoFisher Scientific) and visualized using a Molecular Imager® (ChemiDoc XRS+, Bio-Rad).

#### Indirect Immunohistochemistry

Fish embryos were washed in PBS before fixation in either 10 % buffered formalin or 4 % buffered paraformaldehyde (PF), pH 7.5, for 24-48 h. The fixed tissues were dehydrated through a graded series of ethanol to absolute xylene and embedded in paraffin blocks (Department of Pathology, University Hospital, Univ. Bergen). Sections of the paraffin-embedded embryos (3-5 µm) were attached to poly-L-lysine-coated glass slides (Sigma, St. Louis, Mo) dried at 55 °C for 1 h and de-waxed twice in xylene (10 min each). Sections immersed twice-in absolute ethanol (5 min) and lower ethanol concentration ending to Tris-buffer, pH 7.5. Antigen retrieval of formalin-fixed paraffin-embedded (FFPE) followed the heat-induced antigen retrieval protocols (Cattoretti and Suurmeijer, 1995; Shi et al., 1995; Hayashi, 2011), with subsequent rinsing in TTBS buffer. Sections were incubated in blocking solution using swine or goat normal serum (Dako) diluted in TBS buffer. Immunofluorescence experiments were performed using diluted purified IgG fractions in blocking solution (1:300 and 1:500) with incubation in a humidity chamber overnight, 4 °C. Next, the sections were washed twice in TTBS buffer before incubation in a diluted solution of Fluorescein (FITC) conjugated F (ab’)₂ fragment goat anti-rabbit IgG (Dako) or Alexa Fluor 647 conjugated F(ab)_2_ fragment swine anti-rabbit secondary antibody (Molecular Probes). After washing in Tris-buffer, sections were mounted (Vectashield®, containing DAPI) and visualized, either using TCL-Laser scanning confocal microscope (Sars Centre), or Leica DM-microscope or Leica DMLB fluorescence microscope (MIC-core facility, UiB), using appropriate optical filter sets.

#### Whole mount *Oikopleura* larvae immunohistochemistry

*Oikopleura* larvae were collected after hatching on ice and fixed immediately in 4 % PF (0.1 M MOPS, 0.5 M NaCl) for 2 h. Larvae were washed in PBS (pH 7.0) containing 0.1 % Tween-20 and 2 mM EDTA before incubating in a blocking solution (3 % acetylated bovine serum albumin (Sigma, MO), 0.1 % Tween-20, in PBS) overnight, at 4 °C. Immunofluorescence experiments applied solution of the diluted purified IgG fractions in blocking solution (1:300) with incubation in a humidity chamber overnight, 4 °C. Next, larvae washed and post-fixed in 4 % PF in the presence of 0.1 M MOPS and 0.5 M NaCl overnight, at 4°C. Repeated washes were performed before incubation with secondary antibody (FITC-labelled swine anti-rabbit F (ab’)_2_ (Dako) or Alexa Fluor 647-labelled goat anti-rabbit IgG (Molecular Probes) diluted in blocking solution (1: 500) overnight at 4 °C. Subsequently, specimens were extensively washed and mounted on depression slides (VWR) using VectaShield (Vector Laboratories, CA). Hematoxylin-Eosin and PAS stains were used as described (Bancroft and Stevens, 1996).

#### Total RNA preparations

The salmon total RNA sample (2.0 μg) was extracted from embryos (370dd) inside the 10 ml tube using a Polytron ® PT 1200 E homogenizer in the presence of 2 ml Trizol solution following instructions (Invitrogen, Life Technologies). The zebrafish total RNA samples (1.5 μg) were extracted from embryos at different developmental stages by homogenization in presence of 1 ml Trizol solution inside nonstick, RNase-free microfuge tubes (2 ml tubes, ThermoFisher Scientific). The quantity of the RNA samples was determined spectrophotometrically using the Epoch instrument, while the RNA sample integrity and purity was assessed on the formaldehyde agarose gel Ethidium bromide-stained. The enriched poly (A+) fraction of the RNA was affinity-purified from 5-10 µg of total RNA sample using Superparamagnetic Dynabeads®, coupled to oligo-(dT)25 following the instructions (Invitrogen). The total RNA (2 μg) was reverse transcribed using SuperScript II ribonuclease H-RT (Invitrogen) in 20 μl reactions following instructions.

#### RT PCR

First-strand cDNA was synthesized from a total RNA sample (2 μg) using SuperScript™ II ribonuclease H-RT (Invitrogen) in 20 µl volume in the presence of gene-specific (GS) primer. RNA complementary to the cDNA was removed by adding 1 μl (2 units) of E. coli RNase H with incubation at 37 °C for 20 min.

**PCR amplification** was performed in 50 µl reaction volume using Platinum ^TM^ *Taq* DNA polymerase (5U / μl), DNA-free (Invitrogen) in mixture with other provided PCR components. Samples of cDNA (2 μl, 1:10 dilution) served as a template DNA in combination with GS primers.

**Salmon Leukolectin (sLL)** gene sequence (620 bp) was amplified using following LL-GS primers LL/F 5′-TACGGACACAGGTCGAATCCCCTACTACC-3’ and LL/R 5′-CAGAGAAGAGGCTAATGTGTGCAC-3′. β-actin gene sequence was amplified using β-actin GS primers F 5′-CCCTTTTCCAACCATCCTT-3′ and R 5′-TGGGCCAGATTCATCGTATTCT-3′ as a positive control for equal amounts of cDNA load. The PCR cycling profile involved 35 cycles where each cycle consisted of denaturation at 95 °C (25 sec), annealing at 55 °C (30 sec), and extension at 70 °C (30 sec). Coding gene sequence of **salmon Low Choriolytic enzyme (sLCE) and zebrafish HE 1 gene** (Accession # NM 213635.2; Gene ID: 407971 he1b) were amplified and sequenced as described (Miftari et al., 2022).

Amplified PCR products were resolved by gel electrophoresis on a 1.2 % Ethidium bromide-stained agarose gels (1x TBE-buffer). For sequencing, Platinum *Taq*-amplified PCR products were ligated into Invitrogen pCR4-TOPO TA Vector (Invitrogen TOPO TA Cloning Kits for Sequencing) followed by transformation of the provided One Shot Chemically Competent cells. Subsequently, purified plasmid DNA was sequencing in both directions using primers M13F or M13R at Core Sequencing Facility, University of Bergen.

**For *in vitro* transcription** the insert was excised from pCR4-TOPO TA Vector using flanking EcoR1 site and sub cloned into the multiple cloning sites (MCS) of the pGEM T-Easy vector (Promega) by ligation. Chemically competent cells were transformed, and cultivated at 37 °C, overnight, with shaking. To generate LL-anti-sense or sense riboprobe, ∼1 µg plasmid DNA was linearized (to generate 5’overhang) using a restriction enzyme such as *Spe*1 or *Not* I respectively. To construct β-actin anti-sense or sense riboprobes, plasmid DNA was linearized using restriction enzyme Apo I or *Spe*1, respectively. Riboprobes specific for sLCE or HE1 were prepared from *Nco*I- and Spe1-linearized plasmid DNA. DIG-RNA labelling kit (Sp6/T7) (Roche Diagnostics) was used for DIG or Fluorescein RNA labelling following instructions. Sp6 RNA polymerase synthesize antisense riboprobe using *Spe*1 linearized plasmid DNA, while T7 RNA polymerase synthesize sense riboprobe using *NotI* linearized plasmid. *In vitro* transcription reactions were ended by adding 18 mM EDTA and the generated riboprobes were precipitated in a solution of 5 µg glycogen (5 mg/ml), 80 mM LiCl and 70 % ice-cold ethanol, and incubated at −80 °C for 2 h before storage. The DIG-labelled RNA probes were centrifuged at 13,000 rpm for 30 min at 4 °C washed with 70 % in DEPC-treated water, centrifuged again for 15 min, air-dried for 20 min, and dissolved in 20 µl DEPC-water.

**Northern blot analysis procedures (**Streit et al., 2009) used aliquots of 1 µg total RNA sample and 0.5 µg of poly(A+) RNA, separated by agarose-formaldehyde gel electrophoresis at a constant voltage of 20-30 Volts (15-18 mA) overnight. Subsequently, a capillary blotting system transferred the RNA samples to a positively charged nitrocellulose membrane (Amersham). The RNA blots are immobilized on the nitrocellulose membrane through covalent linkage using transilluminator UV light (254 nm or 302 nm) for 2 min. Membranes were pre-hybridized in DIG Easy Hybridization buffer (Roche) at 55 °C for 2 h, then hybridized at 55 °C for 16 h with one µg/ml riboprobe. The washing was performed twice at low stringency in 2x SSC with 0.1 % SDS at RT for 15 min each, and twice at high stringency in 0.1x SSC with 0.1 % SDS at 65 °C (each 20 min). The membranes were incubated in a blocking solution containing maleic acid buffer (pH 7.5) and 1 % blocking reagent (Roche). Subsequently, the membranes were incubated with anti–DIG –alkaline phosphatase (AP) conjugated antibody (Roche) diluted 1: 10.000 in blocking solution at RT, for 1 h. After washing (0.1 M maleic acid buffer pH 7.5, 0.15 M NaCl and 0.5 % Tween 20), the membrane was equilibrated for 2 min in detection buffer (0.1 M Tris/HCl, 0.1 M NaCl, pH 9.5 at RT). Hybrid probe-targets were visualized by chemiluminescent assays using Roche CSPD chemiluminescent substrate as a substrate. Blots were exposed (5-30 min) to Kodak X-ray film.

***Single labelled in situ* hybridization (***ISH*) procedure on paraffin-sections was performed (Barthel and Raymond, 1993). Briefly, tissue sections were treated with 5 µg/ml proteinase K (Merck) in 100 mM Tris-HCl buffer (pH 8.0) containing 50 mM EDTA at 37 °C for 5 min. Next, sections were post-fixed in 4 % PF in 0.1 M PBS (pH 7.4) at RT (30 min) and acetylated in freshly prepared 0.1 M Triethanolamine-HCl (pH 8.0) with 0.25 % acetic anhydride at RT for 10 min. Approximately 200-400 ng/μl DIG-labelled antisense riboprobe was added in hybridization buffer, and sections were incubated for 16 h at 55 or 62 °C in a humid chamber covered with hybridization cover slip (HybriSlip, Bio-Labs). Next, sections were washed twice for 15 min in 50 % formamide in 5x SSC at 65 °C, 2x 15 min with 5x SSC at RT, twice for 15 min in 0.2x SSC at RT, and once for 5 min in 0.2x SSC at RT. Finally, as a control, many tissue sections follow hybridization protocol with a DIG-labelled sense riboprobe.

**Dual label *in situ* hybridisation** (ISH) was performed on paraffin-sections, as described (Nüsslein-Volhard and Dahm, 2002). The probes were hybridized at 58 °C, or at 62 °C with similar results.

#### Library screening

Salmon library CHORI-214, Seg-1. (Filter set-007193) was screened using digoxigenin-11-dUTP (DIG) labelled LL-specific cDNA probe. Probe was prepared by PCR amplification using forward primers LL/F 5′-TACGGACACAGGTCGAATCCCCTACTACC-3′ and reverse primer LL/R 5′-ACAGAGAAGAGGCTAATGTGTGCAC-3′ in the presence of DIG (Roche). PCR-products were resolved in a 1.2 % agarose Ethidium bromide-stained gel. DIG-labelled cDNA probe was incubated at 95 °C for 10 min and immediately placed in the ice for 5 min before adding to the hybridization buffer (5x SSC, 50 % formamide, 0.02 % SDS, 2 % blocking agent (Roche), DEPC-treated water). Hybridization was carried out in hybridization tubes overnight at 55 °C. The post-hybridization protocol included two washes in a low stringency washing buffer (2x SSC, 0.1 % SDS at RT, 15 min each) and two washes at a high stringency washing buffer (0.1x SSC, 0.1 % SDS at 65 °C, 20 min each). Next, filters were incubated in a blocking solution containing maleic acid buffer (pH 7.5), 1 % blocking reagent before incubation in a blocking solution with anti-DIG alkaline phosphatase (AP) conjugated antibody (Roche) diluted 1: 10.000 at RT, for 1 h. After washing with 0.1 M maleic acid buffer (pH 7.5) containing 0.15 M NaCl and 0.5 % Tween 20, the membranes were equilibrated for 2 min in detection buffer (0.1 M Tris-HCl, 0.1 M NaCl, pH 9.5) at RT. Filters were subjected to a chemiluminescent assay using CSPD (Roche) as a substrate and exposed to Kodak X-ray film, with 3-15 exposure times. The positive duplicate spots (clones (L-7/257 and A-12/144) were plated and incubated on agar plates at 37 °C overnight. Three clones were selected for cultivation overnight in a 5 ml Luria Bertani (LB-) medium under shaking at 37 °C in the presence of 20 µg/ml chloramphenicol. Plasmid DNA purification used an ultrapure DNA purification kit (Qiagen). BAC plasmid DNA was digested with *Not* I before analysis by pulse-field gel electrophoresis (Osoegawa et al., 1998). Positive clones were verified by PCR amplification using purified BAC plasmid DNA as a template and gene-specific primers. The PCR amplification profile included initial incubation at 95 °C for 3 min, followed by 35 cycles at 95 °C for 30 sec, annealing at 54 °C for 30 sec, and extension at 72 °C for 30 sec with a final extension at 72 °C for 10 min. PCR-generated amplicons were analyzed on a 1.5 % Ethidium-bromide agarose gel.

#### Shotgun sequencing of LL-specific BAC clones

MRW, Berlin, Germany, carried out sequencing of clones. The vector inserts were sequenced to an estimated 2.4fold redundancy. From the 180 reads, 33 and 22 contigs were established using bioinformatics from the two BAC clones. The analysis identified exons and introns by program searches and manual inspections. The entire LL gene was identified from overlapping contigs. Here, L7 gene is reported.

**Zebrafish Leukolectin (zLL**) cDNA-sequence was amplified using sLL-specific primers (Table 1). The PCR mix (20 μl) contained 2 μl of cDNA template, 0.4 μM of primers (LL-F1, LL-F2, LL-R1, LL-R2, and LL-R3), 1x PCR buffer, 0.5 U Taq DNA polymerase (Takara), and 0.2 mM of dNTPs. PCR amplification used 32 cycles of 94 °C for 30 sec, annealing at 58 °C for 45 sec, extension at 72 °C for 90 sec, with final extension at 72 °C for 10 min. The reaction mixture was analysed by 1.2 % agarose-TBE gel with 0.5 µg/ml Ethidium bromide in 1x TBE buffer (pH 8.3).

**Table 1:**
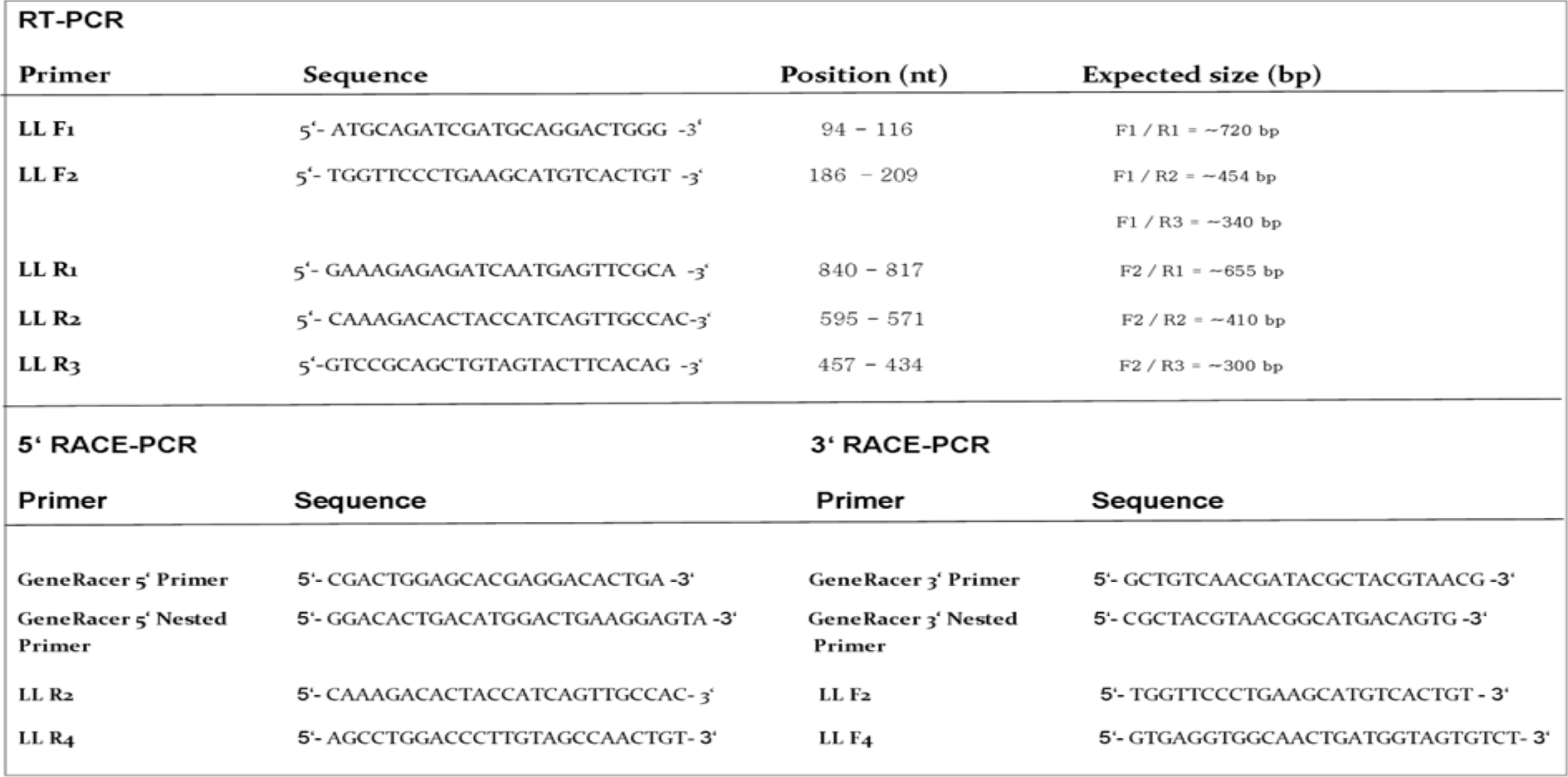
Primers used for zebrafish LL amplification

**zLL 5’- and 3’- RACE PCR.** zLL 5’- and 3’- RACE PCRs were performed using the GeneRacer core kit (Invitrogen). Total RNA (2 µg) was used to synthesize the first-strand cDNA using Superscript III Reverse transcriptase. The first round of 5’- and 3’-RACE PCR amplification was performed by applying 5’- or 3’- GeneRacer primer and reverse or forward LL-GS primer LL-R2 and LL-F3 (Table 1). RACE PCR applied touch down PCR, where the first step consisted of five cycles (94 °C for 30 sec; 72 °C for 60 sec), and second step of five cycles (94 °C for 30 sec; 70 °C for 60 sec), and the third step of 30 cycles (94 °C, 30 sec, 68 °C, 45 sec, 72 °C, 1 min) with final extension step at 72 °C, 10 min). Templates for nested PCR used PCR product from first-round RACE PCR diluted 1:25 in destilled water. Amplification included 25 cycles (94 °C for 30 sec, 65 °C for 30 sec, and 68 °C for 1 min), with the final extension step at 72 °C for 10 min. PCR products were purified using PCR purification kit (Promega, WI, US), and ligated into PCR II Topo-vector (Invitrogen) for transformation of JM109 high-efficiency competent E. coli (Invitrogen). Recombinant bacteria were identified by blue/white screening on LB-agar plates. Plasmids containing the inserts were purified using Promega purification kit and sequenced with M13 primers. After sequencing, the putative zLL-protein domain-sequences were analysed by SMART-program (http://smart.embl-heidelberg.de/). Further molecular characteristics were determined using ProtParam (http://web.expasy.or/protparam/). Homology searches in GenBank database used BLAST network server. Multiple sequence alignments used the same server.

**Zebrafish genomic DNA** was extracted from 24 hpf embryos, as described (Westerfield, 2007). Extracted DNA was resuspended in 5 ml of 10 mM Tris pH 8.5, 5 mM EDTA, and gently mixed at 37 °C. Concentration was determined spectrophotometrically at OD 260 nm. Diluted zDNA samples served as template for PCR-amplification, using forward primers designed from contig (Fig. S 5) in combination with reverse primers (Table 1). PCR-amplification cycling profile was as described above for salmon LL-gene amplification.

**Construction of DIG-labelled zLL-riboprobes for** *ISH* **experiments** used yhese LL-primers: LL-F1 and LL-R1 (Table 1) giving a probe of around 670 nt. A second probe was generated using LL-F2, and LL-R2 giving a probe of around half the size (see Table1). Both probes gave the same results. Digoxygenin (DIG) RNA labelling was performed using DIG-labelling kit (Roche) following instructions. Sp6 (or T7) RNA polymerases were used for transcription of antisense or sense probes from respectively *Spe*1, *Not* I (or *XhoI*)-digested plasmid. *In vitro* transcription reactions were stopped by 18 mM EDTA. Products were precipitated and incubated at −80 °C for 2 h. The prepared DIG-labelled RNA probe was dissolved in 20 µl DEPC-treated water.

**Whole-mount zebrafish *ISH*** was performed as described (**Seo et al., 1998**). Chromogen substrate NBT/BCIP (Roche) was used to detect DIG-labelled riboprobe. After the whole-mount *ISH* procedure with a single riboprobe, embryos were paraffin-embedded and serially sectioned at 3-5 µm (Leica microtome), before attachment to poly-L-lysine-coated glass slides (Sigma, St. Louis, Mo). Sections were examined by light microscopy, and images were taken as described below.

#### Combined whole-mount *ISH* and immunohistochemistry in zebrafish embryos

This procedure was performed as described (Streit, A. and Stern, C. D., 2001; Nüsslein-Vollhard and Dahm, 2002). In short, specific fluorescein-derivatized riboprobes for LL or for HE1, and visualized by AP-conjugated antibodies, and revealed by Fast Red chromogen (red colour). Antigens (for HE1-, or LL-) proteins were detected using 10 mg tablets of DAB (Dako Cytomation) diluted in 5 ml Tris-buffer (brown colour).

#### Imaging

Leica TCS SP5 confocal microscope equipped with a 5.0 RTV camera (Q Imaging) used for capturing the fluorescent images. Leica TCL laser scanning confocal microscope (for salmon and *oikopleura*) was equipped with 40x and 60x oil objectives (numerical aperture 1.25 and 1.40, respectively). Image stacks of about 1 µm optical sections acquired using Leica powerScan software. Leica DM-microscope or Leica DMLB fluorescence microscope equipped with a 40x oil immersion objective (numerical aperture 1.00) used for capturing the images from *ISH* experiment. Whole-mount *in situ* hybridized images employed Leica MZ 16A microscope, numerical aperture 1.00, equipped with Leica camera 1044626, and Nikon camera digital sight DS-U1. Images are processed with the photo program ACT-2U, and montage images are made with the auto montage program. Light microscopy used a Leica DMLB microscope. Adobe CS2 Photoshop (San Jose, CA) was used for image contrast and brightness adjustment.

## Acknowledgements

This work was supported by German Academic Exchange Service (DAAD#324, 2000; to MHM); Norwegian Research Council Fellowship (2001-2002, to MHM); Norwegian Research Council (Regional Research Funds (RFF-grant 2012-2016). Counseling from profs. H.J. Seitz, Univ. of Hamburg, Germany, K. Pavelic, Univ. of Zagreb, Croatia, and the late S. Roseman, The Johns Hopkins University, USA, was invaluable. Support from Arild Rasmussen and Reidar Holmefjord (AS Bolaks), and from Thorleif Thormodsen (Alginor ASA) was crucial. Assistance from GC Rieber ASA (T. Eide), J-M. Boquet (Sars Centre), B. Nordanger (Dept. of Pathology, Univ. of Bergen) and J. Fagertun (Fusa) is gratefully recognized. Technical support from Molecular Imaging Center (MIC) and from the Core Sequencing Facility at the University of Bergen is acknowledged.

## HIGHLIGHTS

- Lectocytes synthesize leukolectins-proteins (LL) for exocrine secretion *in ovo*
- Lectocytes are defined as special embryonic peridermal mucous cells
- Lectocytes are among the earliest functional embryonic cells
- LL is involved in first-line, dermal innate immunity defence
- The LL-gene exhibits particular upstream transcription regulation
- Conserved LL-proteins with 5 TECPR-domains belong to the tectonin-family

## SUPPLEMENTARY FIGURES

**Fig. S1:**
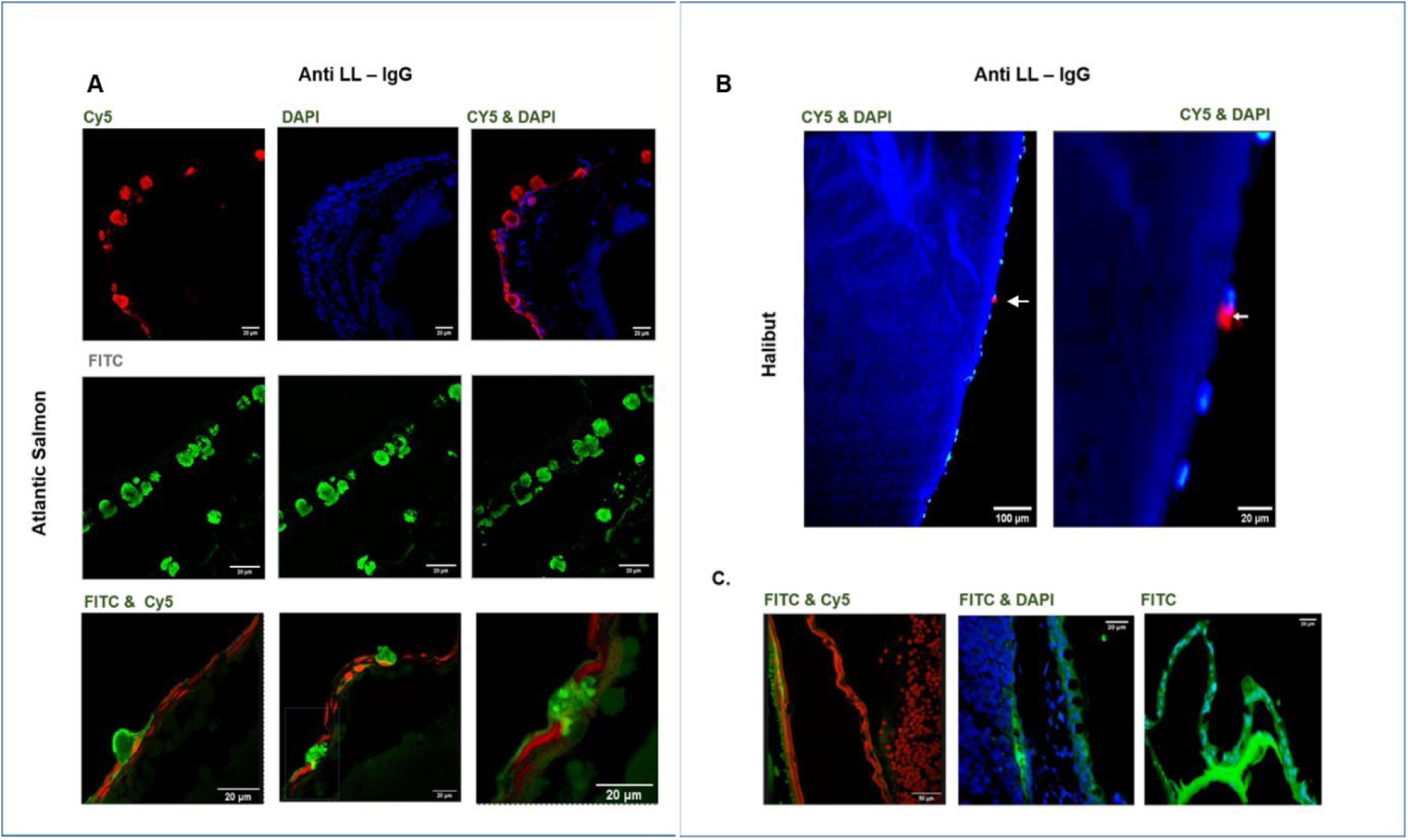
Immunofluorescent analysis of salmon and halibut embryonic sections. Paraffin-sections (3-5 µ) of (4 % paraformaldehyde) fixed embryos, processed as in Fig. 1. **A:** Confocal microscopy of immuno-stained salmon sections. Top panel: Right image presents merged view of left (Cy5) and middle images (DAPI), showing immuno-stained cells in red colour. Middle panel: View of immuno-stained cells (green) without counterstain. Notice prominent granulated features of lectocytes. Lower panel; Left and middle merged images show position of immune-stained lectocytes relative to the peridermal surface. Notice granulated appearance of lectocytic cytoplasm (right image). Nuclei are propidium-stained (viewed in Cy5). **B**. Fluorescent microscopy of immuno-stained halibut sections. Left image shows merged view of rare immuno-stained cells. Right image: high magnification of lectocyte (white arrow). **C**. Negative control. Images of salmon (left and middle) and rainbow trout (right) sections after processed without primary antibody. Filters and scale bars indicated.

**Fig. S2.**
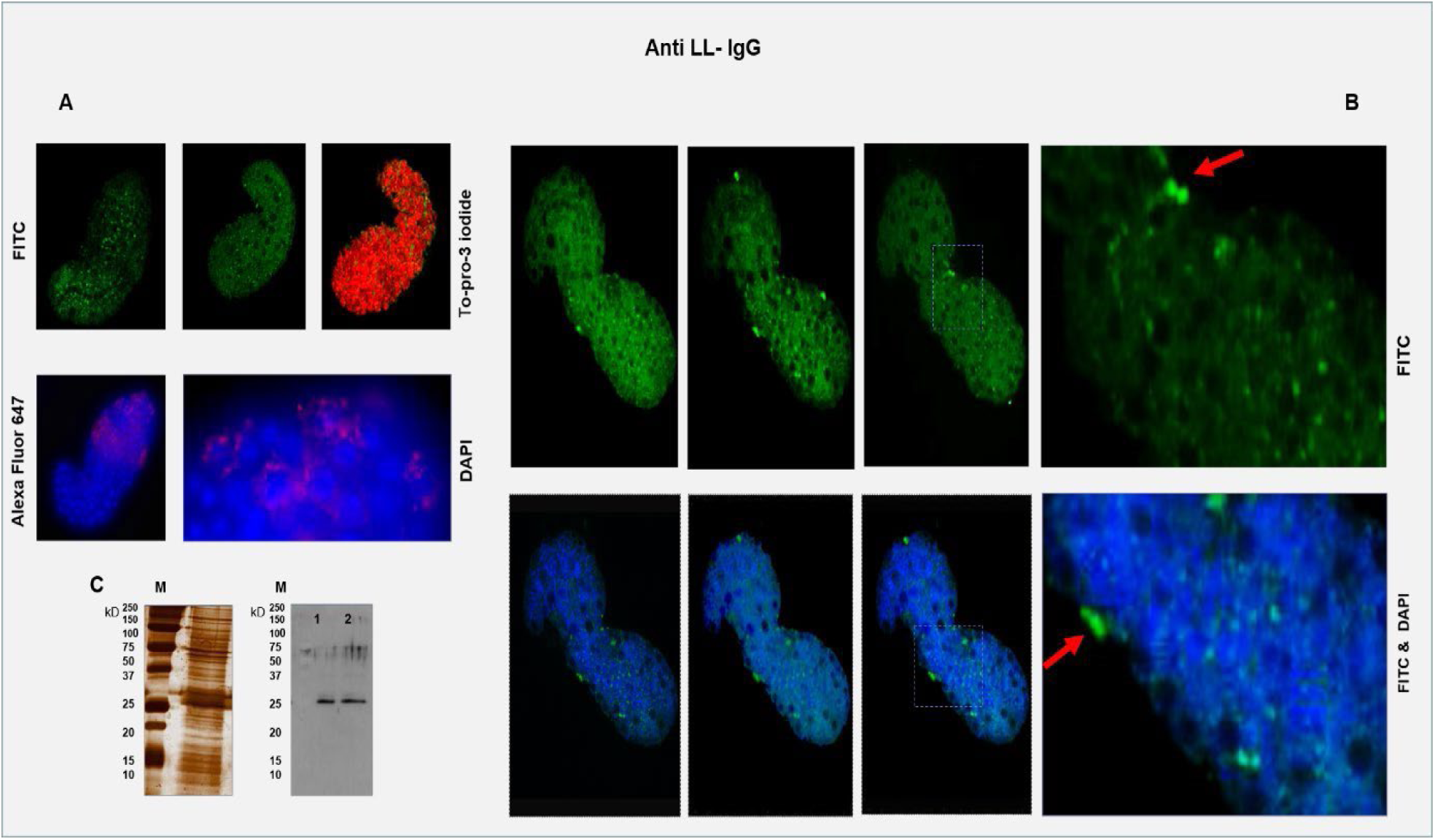
Immuno-stained cells in whole mount *Oikopleura dioca* larvae using LL-IgG. **A**: Larvae at 5 hpf **Upper panel:** Optical section images from confocal microscopy: <u>Left</u> <u>image</u>: LL-immuno-reactive cells (green) are rounded and dispersed with moderate staining intensity. Nuclei not counterstained. <u>Middle image:</u> Immuno-reactive cells. <u>Right image:</u> Composite reconstructions of whole larva from serial optical sections. Nuclei (red) stained by To-pro-3 iodide, while (green) immuno-reactive cells are detected by FITC-conjugated secondary antibody. Notice minority of non-clustered LL-immunoreactive cells throughout the larva. **Lower panel:** Fluorescence microscopy of larva. <u>Left image:</u> Immuno-reactive cells (red) are prominent in the head region, exhibiting varying staining intensities. Merged image of views from Alexa Fluor 647 (red) and DAPI (blue, nuclei stain) filters. <u>Right image:</u> High magnification from left image. Clusters of immuno-negative (blue) cells often appear with interspersed multiple immuno-reactive cells. **B**: Larvae around 6 hpf. Confocal microscopy. **Upper panel** show images of (green) immuno-reactive cells in selected optical sections (FITC-filter): <u>Left three images</u> show immuno-reactive cells (bright green) located inside the soma as smaller and weakly stained. External immuno-reactive cells are larger and intensively stained. <u>Right image</u>: higher magnification (from box in previous image). All images are without nuclear stain. **Lower panel** show such selected optical sections merged with DAPIviews, (except for alternate boxed area). <u>Notice</u> that the minority of highly immuno-stained cells with external location, often seen in clusters (Red arrow). **C:** Left image: Proteins *oikopleura* HF (15 % SDS PAGE) are silver-stained in right lane. Right image: Western blot of *oikopleura* HF-proteins (15 % SDS PAGE), analysed in duplicates as described in Methods. Left lane: Negative control. Two right lanes show HF (collected from 5, respectively from around 6 hpf). A main immuno-reactive protein of around 26 kDa. M: BioRad’s Dual color 161-0374 marker Confirmation of antigen-identity by transcript expression profile is needed to establish the identity of oikopleura’s LL-immunoreactive component. **Abbreviation**: hpf = (hours post-fertilization) <u>Ref</u>**: Nishida, H.** (2008) Development of the appendicularian Oikopleura dioica: culture, genome, and cell lineages. *Dev Growth Differ*. **Supp. 1,** S529-S 556.

**Fig. S3.**
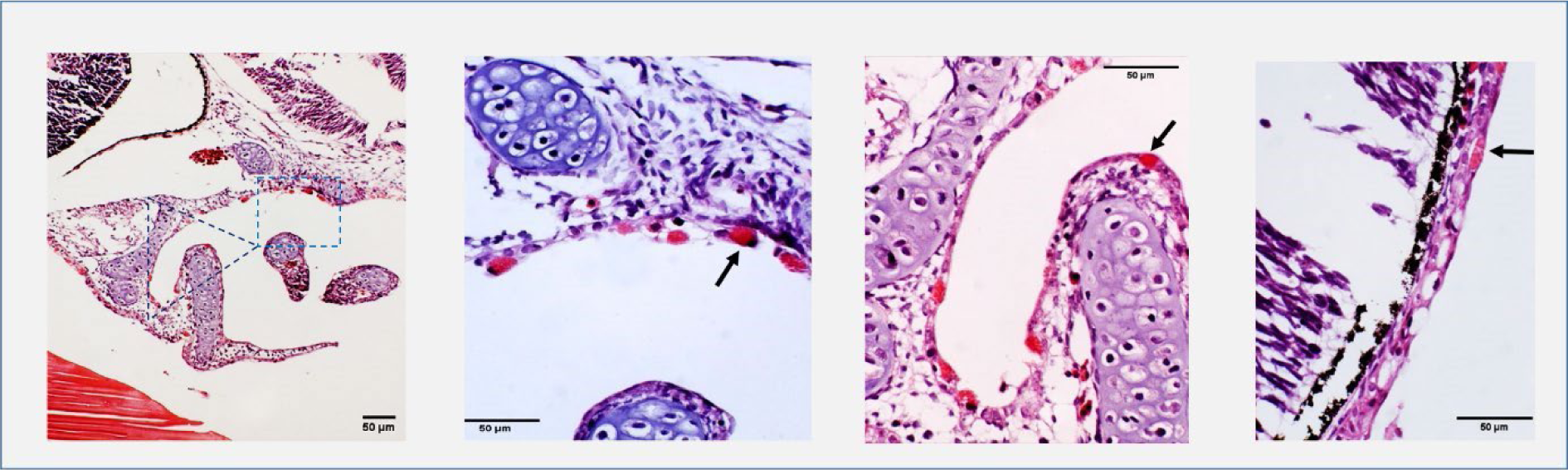
Light microscopy images of Eosin-stained paraffin-sections of Rainbow trout embryos. Left image shows low magnification of embryonic gill-region. Second image shows higher magnification from the Boxed-area in left image. Arrows point to hatching glands stained red by eosin. Third image shows higher magnification of triangled area in left image. Arrow points to Hatching gland located in between two epithelial dermal cell layers. Right image shows view of the embryonic dermis, where the arrow points to a single (red) hatching glands. Notice adjacent empty-looking cells (not stained by eosin) which are more numerous with an external location. Scale bars indicated.

**Fig. S4.**
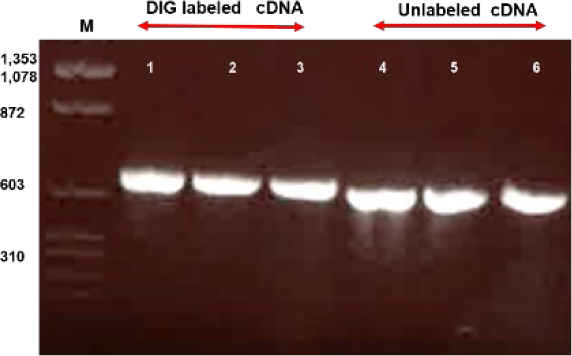
Salmon cDNA-probes synthesized by PCR-amplification. Ethidium bromide (1.2 %) agarose gels showing LL-amplicons. Three left lanes show DIGlabelled amplicons of approximate size 700 bp. Three right lanes show unlabeled amplicons.

**Fig. S5.**
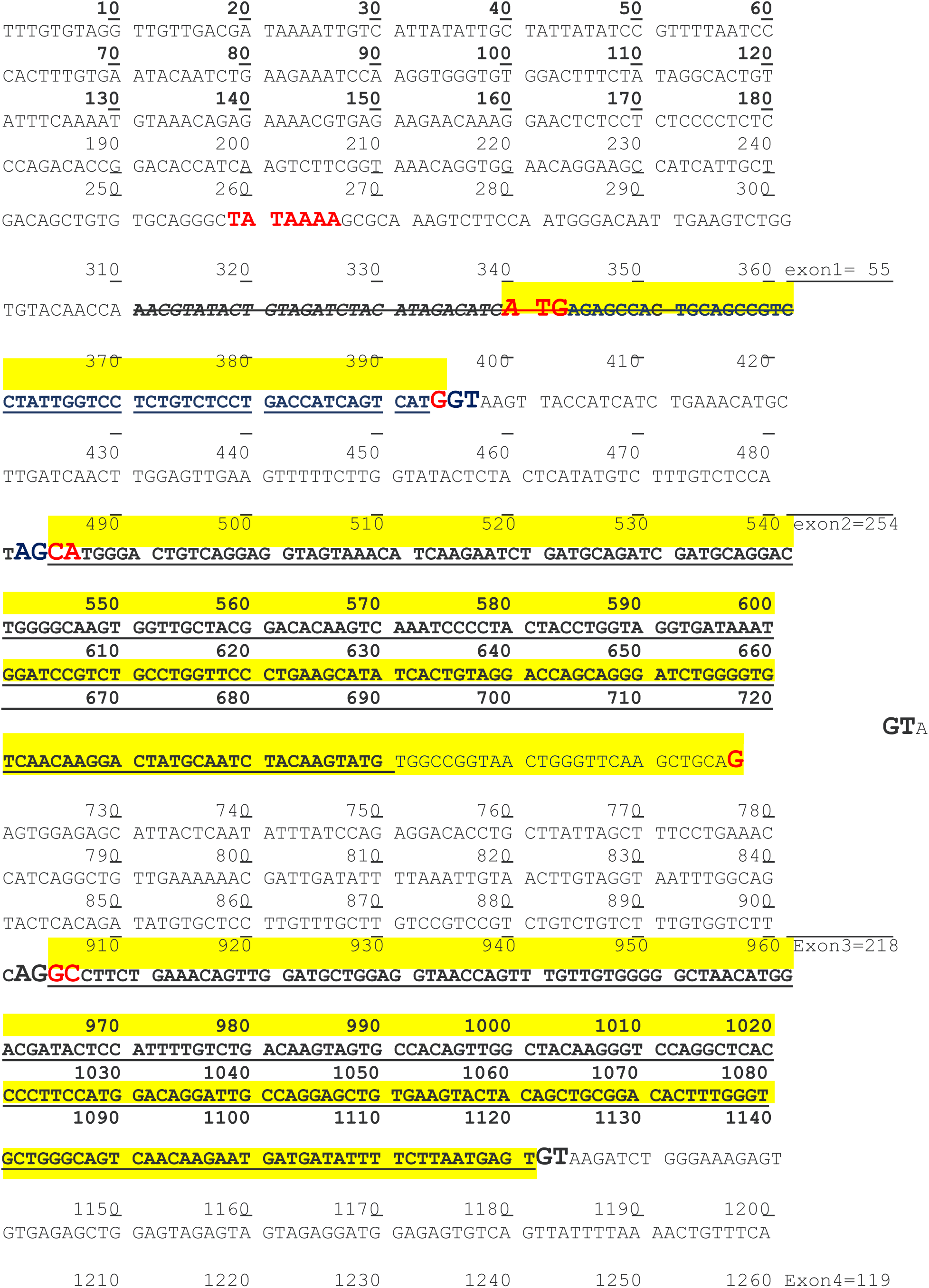
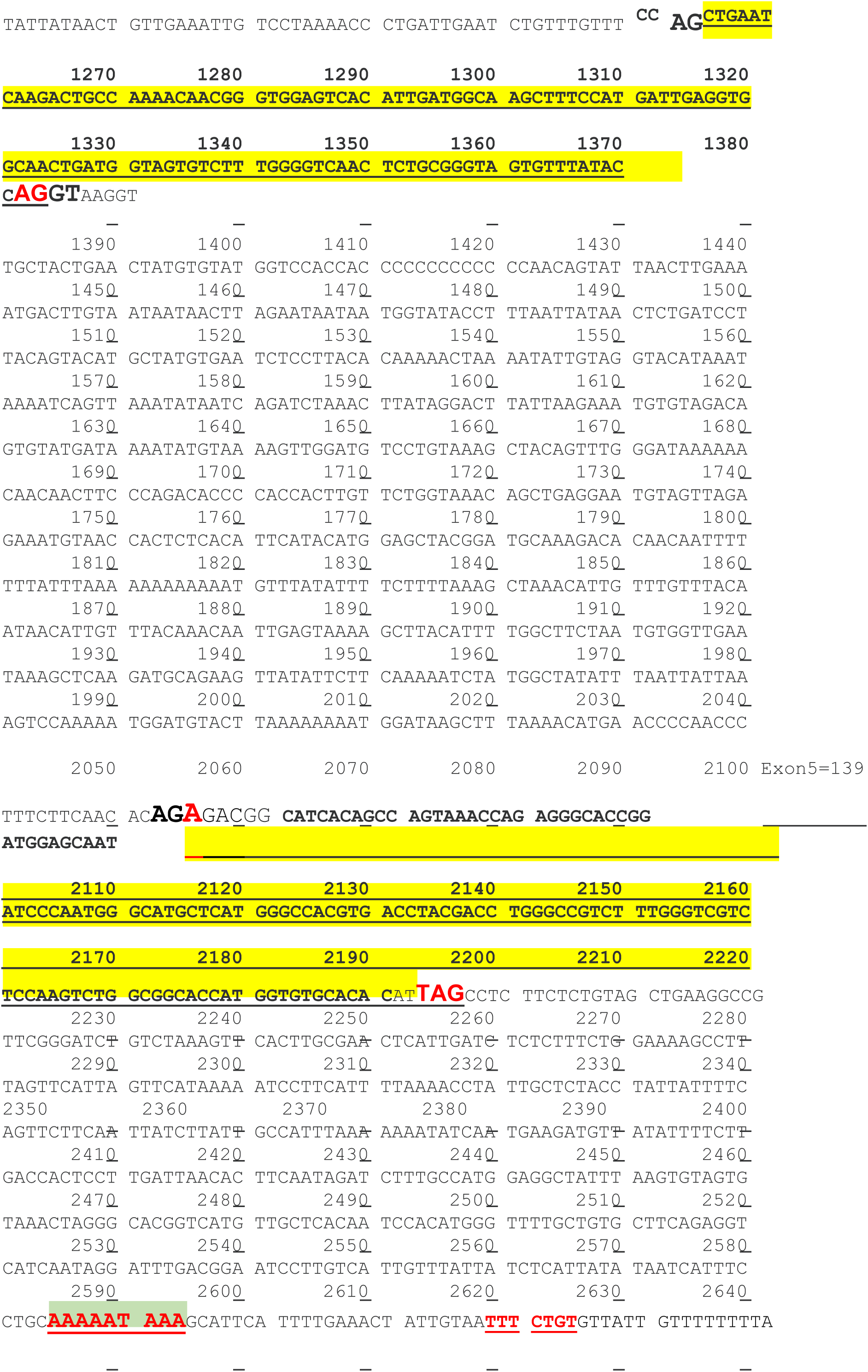
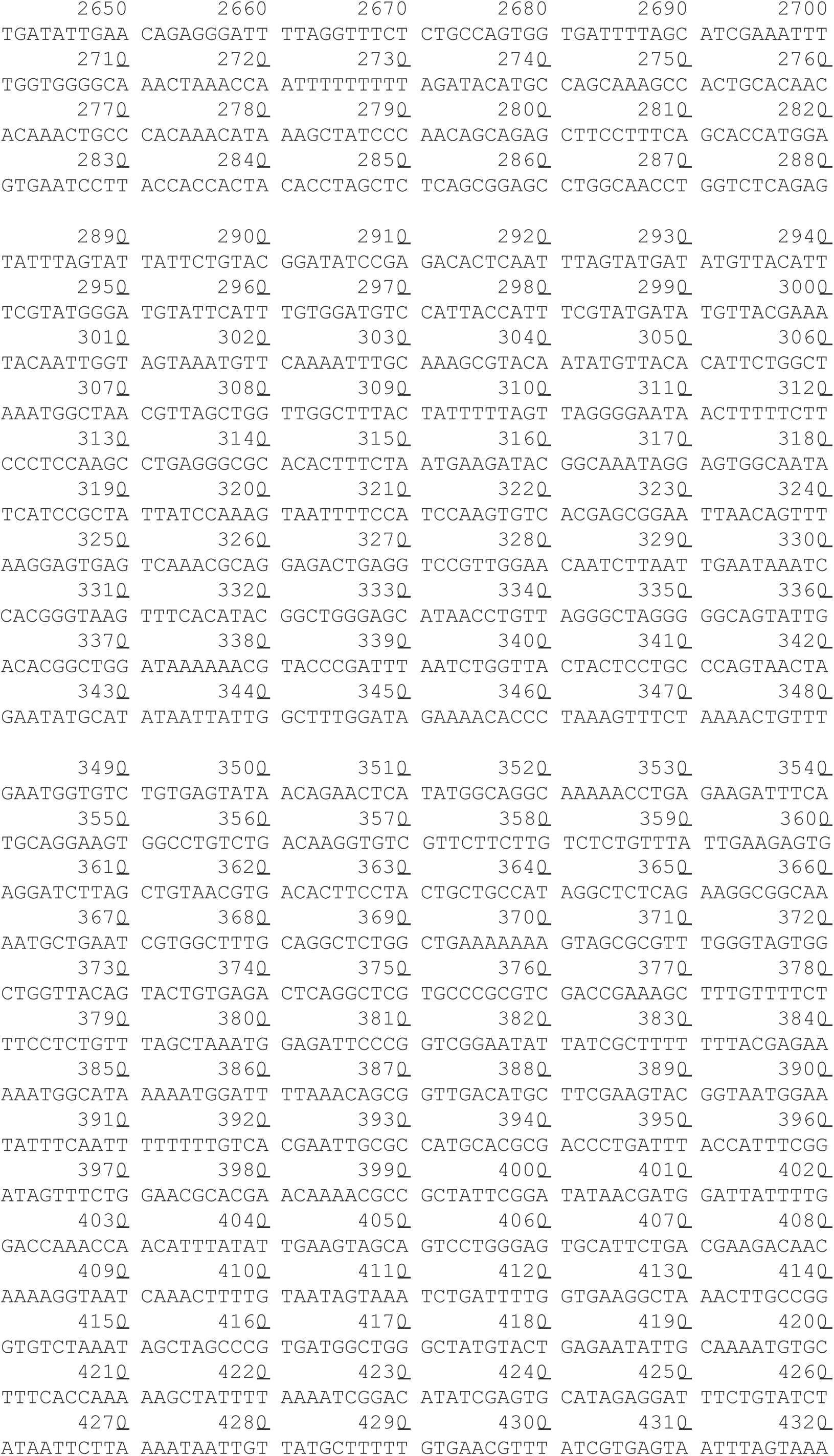
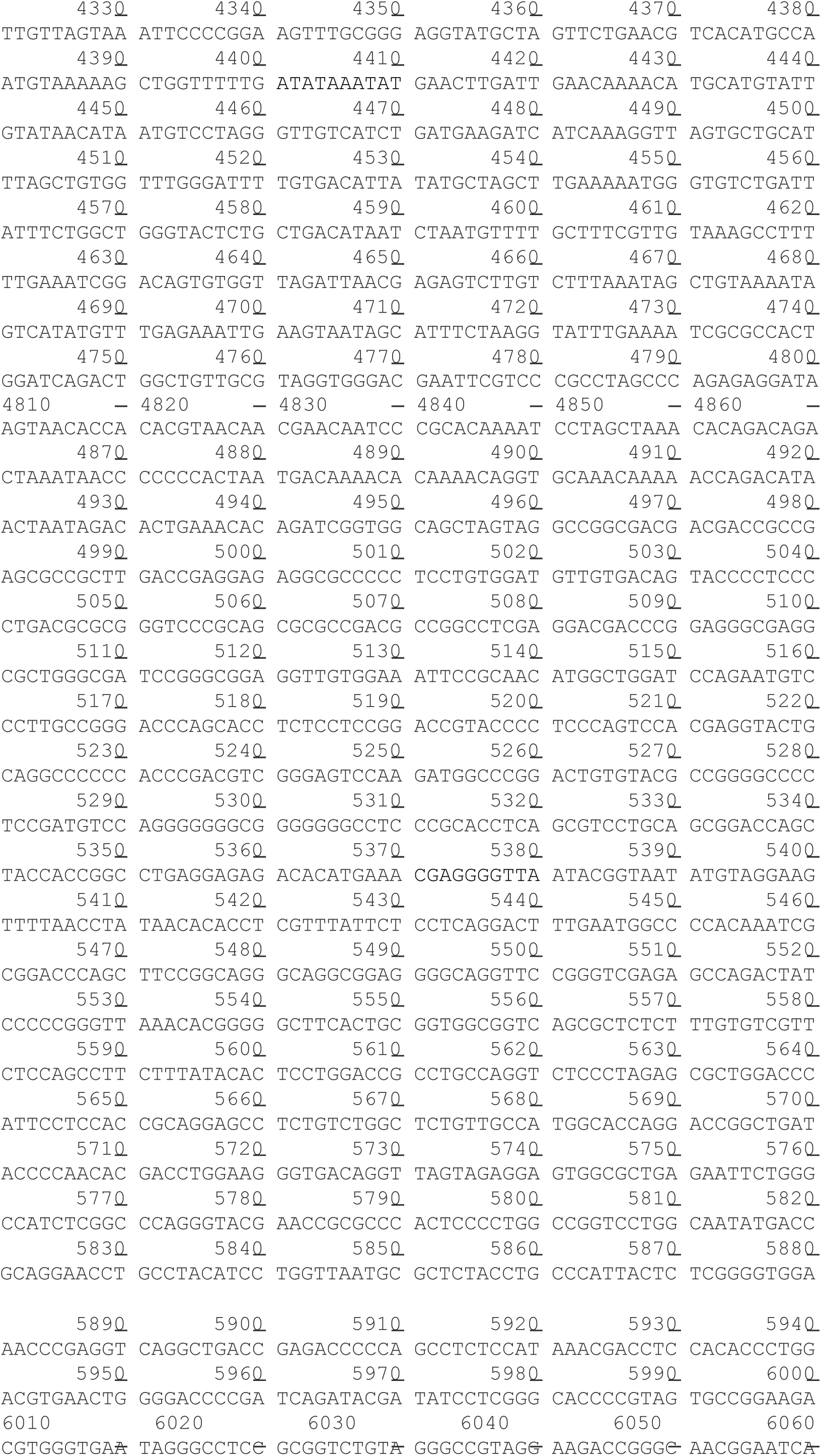
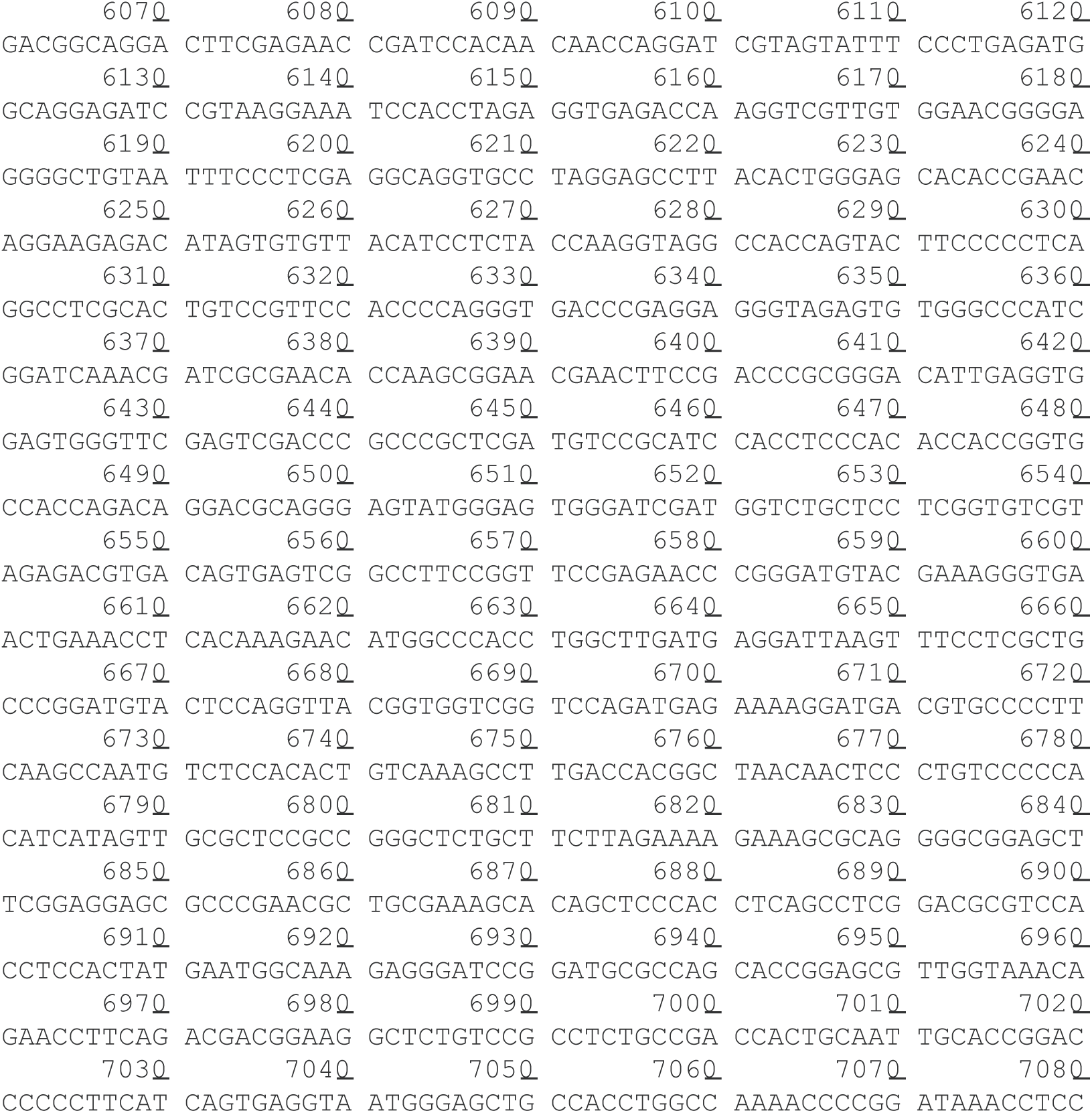
Sequence of DNA-fragment derived from Contig. 14, clone L7. Positions of Exons are indicated along with TATA-box, start codon, splicing acceptor & donors, and polyadenylation signals. For PCR amplification of zebrafish, forward primers from salmon LL upstream region were: F1=5’-CCGGCACCATCAAGTCTTCGGTAAAC-3’; F2= 5’CGTGAGAAGAACAAAGGAACTCTCCTCT-3’. For Reverse primers, see Methods.

**Fig. S6.**
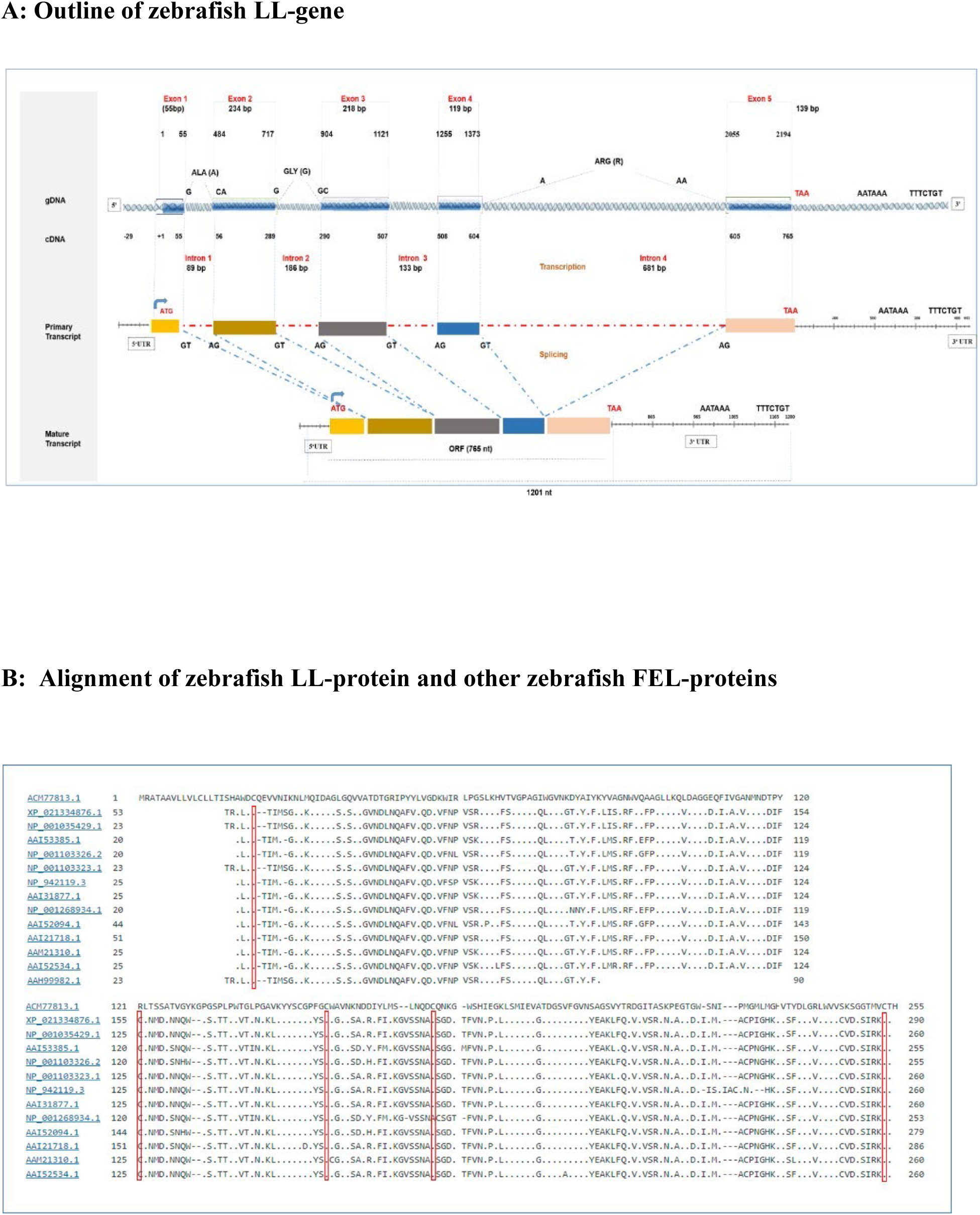
Characteristics of zebrafish LL. **A. Outline of zebrafish LL-gene (zLL-gene).** The amplicons were generated by PCRamplification using zebrafish DNA extracted from embryos at 24 h using different primer combinations, allowing construction of the zLL-gene. The outline demarcates a partial 5’ UTR of some 30 bp, followed by start codon. Splicing sites define sizes of five exons and four introns with split codons for Ala, Gly and Arg. All four splice donors and acceptors are identified as GT and AG, respectively. Positions of stop codons and polyadenylation-site are shown. An ORF of 765 nt is defined, with 3’-UTR of 430 nt. Numbers refer to positions in cDNA. A primary transcript size ̴ 2.300 nt is predicted, with a mature transcript > 1.230 nt. **B. Alignment of putative zebrafish LL-protein and putative zebrafish FEL-proteins. U**sing leukolectins (ACM77813.1:1-255) as query sequence, the following putative zebrafish lectins (presented in Figure, from Top to Bottom) were aligned: XP_021334876.1:53-290 fish-egg lectin-like isoform X1; NP_001035429.1:23-260 Fish-egg lectin-like precursor; AAI53385.1:20-255 Zgc:173443 protein; NP_001103326.2:20-255 Fish-egg lectin-like precursor; NP_001103323.1:23-260 uncharacterized protein LOC100126125 precursor; NP_942119.3:25-260 Fish-egg lectin-like precursor; AAI31877.1:25-260 Zgc:158494 protein; NP_001268934.1:20-253 Fish-egg lectin-like precursor; AAI52094.1:44-279 LOC561528 protein; partial >AAI21718.1:51-286 LOC561528 protein, partial; AAM21310.1:25-260 putative galactose-binding protein; AAI52534.1:25-260 Zgc:158494 protein; and AAH99982.1:23-90 Zgc:109753. While the sequence similarity between leukolectins and these sequences is all cases below 50 %, the alignment indicates common structural elements, such as positions of Cys-residues (marked in red). On the other hand, the exact positions predicted for TECPR-domains in these lectins seem to deviate somewhat, whether four or five domains are predicted. The dash-symbol represents identical residues, while varying AA-residues are represented with one-letter codes.

**Fig. S7.**
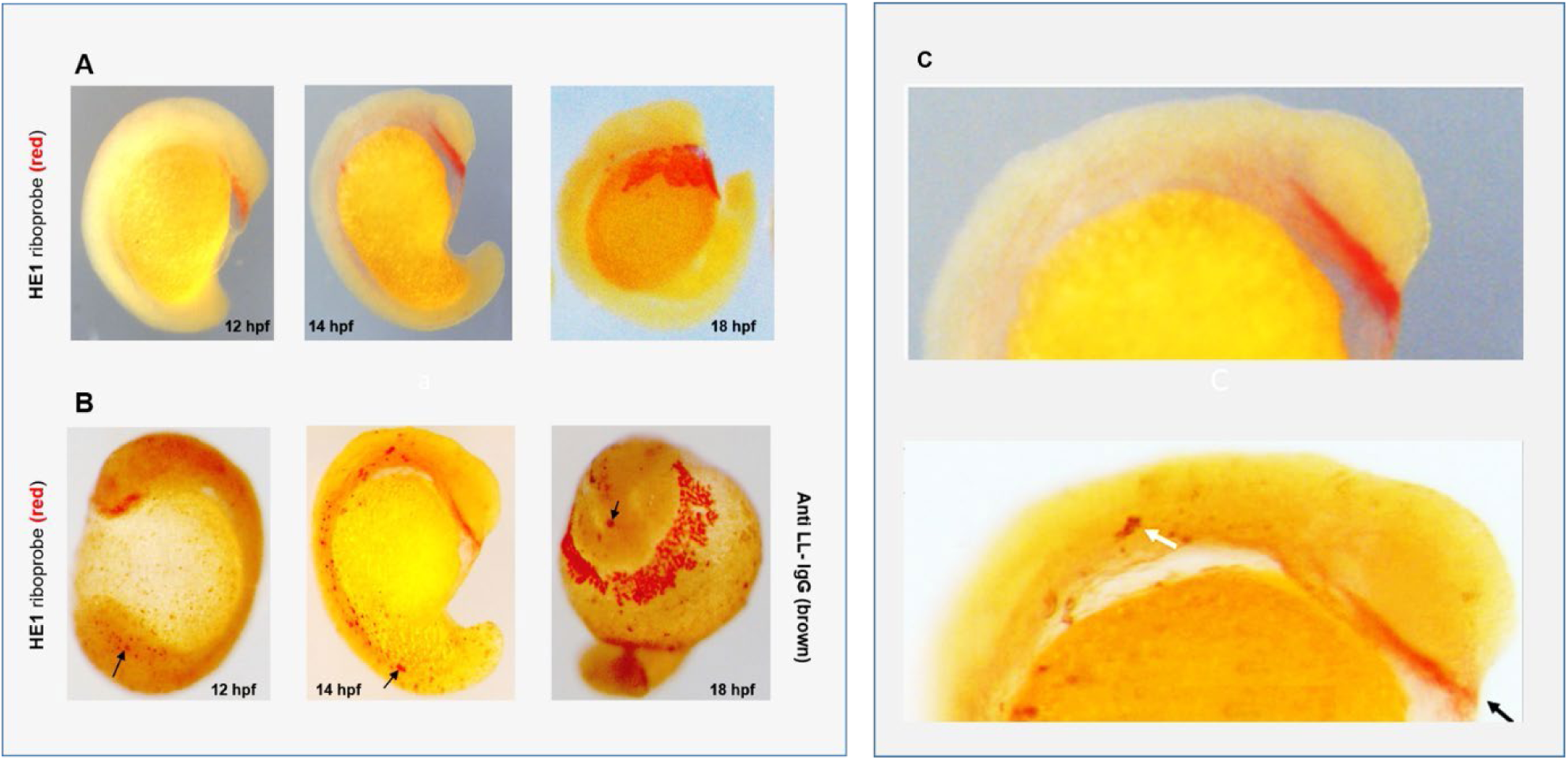
Distinction between zebrafish lectocytes and hatching glands. A. HG-cells revealed by single-labelled, whole-mount zebrafish RNA *in situ* hybridization using DIG-labelled HE-specific anti-sense riboprobe (red colour). Left image: HE1-transcript detected in pre-chordal plate 11 hpf. Middle image: HE1-transcripts are detectable in migrating HG-glands (14 hpf). Right image: HE1-transcripts are detectable at HG-glands in ventral part of the yolk sac (19 hpf). B. Lectocytes in whole mount zebrafish immunohistochemistry, detected by anti-LL antibody (with DAB; brown colour), and their relationship to HG-cells revealed by *in situ* hybridization, using DIG-labelled HE1 specific riboprobe (red colour). Left image: LL immuno-reactive cells are widely dispersed over the embryonic body (11 hpf). Arrow points to LL immuno-reactive cells in caudal part of the embryonic body where HG-glands are *not* present. Middle image: Embryonic body (at 14 hpf) with LL-positive (brown) cells of polymorphic cytology (see arrow). Right image: LL-immuno-positive (brown) cells in cranial part. HG-cells (red) occupy a ventral (necklace) position. C. Higher magnification of A and B. Top image (from A) demonstrates HG-cells in canonical position. Lower image (from B) shows brown lectocytes (white arrow) which are elongated and polymorphic. Black arrow points to (red) hatching glands.

## ABBREVIATIONS

BAC: Bacterial Artificial Chromosome
dd: days x degrees centigrade
hpf: hours post-fertilization
dph: days post-hatching
FEL: Fish Egg Lectin
HE: Hematoxylin-Eosin (stain)
HF: Hatching Fluid
HG: Hatching gland
*ISH*: *In situ* hybridization
LCE: Low Choriolytic enzyme
LL: Leukolectin
PAS: Periodic Acid Schiff (stain)
PRR: Pathogen Recognizing Receptor
PVF: Perivitelline fluid
PVS: Perivitelline space
TECPR: Tectonin protein family Domains

## REFERENCES in METHOD SECTION.

Bancroft, J. D. and Stevens, A. (1996). Theory and Practice of Histological Techniques (4th Ed.) pp. 188–190.New York, Churchill Livingstone.

Barthel, L. K. and Raymond, P. A. (1993). Subcellular localization of α-tubulin and opsin mRNA in the goldfish retina using digoxigenin-labeled cRNR probes detected by alkaline phosphatase and HRP histochemistry. J. Neuroscience Meth. 50, 145–152.

Cattoretti, G. and Suurmeijer, A. J. H. (1995). Antigen unmasking on formalin-fixed, paraffin-embedded tissues using miocrowaves: A Review. Adv. Anat. Pathol. 2, 2–9.

Harlow, E. and Lane, D. (1988) Antibodies, a laboratory manual. Cold Spring Harbor Laboratory. ISBN 0-87969-314-2 (1988).

Hayashi, H., Lewis, A., Hayashi, E., Betenbaugh, M. J. and Su, T-P. (2011). Antigen Retrieval to Improve the Immunocytochemistry Detection of Sigma-1 Receptors and ER Chaperones. Histochem. Cell Biol. 135, 627–637.

Heukeshoven, J. and Dernick. R. (1985). Simplified method for silver staining of proteins in polyacrylamide gels and the mechanism of silver staining. Electrophoresis 6, 103–112.

Laemmli, U.K. (1970). Cleavage of structural proteins during the assembly of the head of bacteriophage T4. Nature 227, 680–685.

Miftari, M.H. (2009). Leukolectins: Novel and Evolutionarily Conserved Proteins with Relevance for Vertebrate Macrophages. *PhD-thesis*, University of Bergen, Norway. ISBN 978-82-308-0842-9.

Miftari, M.H., Bjune, J-I., Rong, C.J., Nilsen, F. and Walther, B.T. (2022). Choriolytic and non-choriolytic proteases during hatching of Atlantic Salmon embryos. Comp. Biochem. Physiol. B. 262, 1–12. doi.org/10.1016/j.cbpb.2022.110776

Nüsslein-Volhard, C. and Dahm, R. (2002). Zebrafish. A practical approach. Oxford, UK: Oxford University Press. ISBN 0 19 963809

Osoegawa, K., Woon, P. Y., Zhao, B., Frengen, E., Tateno, M., Catanese, J. J. and de Jong, P. (1998). An improved Approach for Construction of Bacterial Artificial Chromosome Libraries. Genomics 52, 1–8.

Rong, C.J. (1997). Zonase from *Salmo salar*. Characterization of Endoproteases that Mediate Hatching Transition in the Sexual Life Cycle. PhD Thesis, University of Bergen, Norway. ISBN: 82–7653–010-9

Schoots, A. F., Opstelten, R. J. and Denucé, J. M. (1982). Hatching in the pike (*Esox lucius*): Evidence for a single hatching enzyme and its immunocytochemical localization in specialized hatching gland cells. Dev. Biol. 89, 48–55.

Seo, H-C., Drivenes, Ø., Ellingsen, S. and Fjose, A. (1998). Expression of the murine Six3 gene demarcates the initial eye primordial. Mech. Dev. 73, 45–57.

Shi, S-R, Key, M. E. and Kalra, K. L. (1991). Antigen retrieval in formalin-fixed, paraffin-embedded tissues: an enhancement method for immunohistochemical staining based on microwave oven heating of tissue sections. J. Histochem. Cytochem. 39, 741–748.

Streit, A. and Stern, C. D. (2001). Combined Whole-Mount *in Situ* Hybridization and Immunohistochemistry in Avian Embryos. Methods 23, 339–344.

Streit, S., Michalski, C. W., Erkan, M., Kleeff, J. and Friess, J. H. (2009). Northern blot analysis for detection and quantification of RNA in pancreatic cancer cells and tissues. Nat. Protoc. 4, 37–43. doi.org/10.1038/nprot.2008.216

Towbin, H., Staehelin, T. and Gordon, J. (1979). Electrophoretic transfer of protein from polyacrylamide gels to nitrocellulose sheets; procedures and some applications. Proc. Natl. Acad. Sci. US. 76, 4350–4354.

Westerfield, M. (2007). A guide for the laboratory use of zebrafish (Danio rerio). In *The Zebrafish Book*. 5^th^ Edition, Chapter 9. Eugene, Univ. of Oregon Press.

## REFERENCES

Abrams, E.W. and Mullins, M.C. (2009). Early zebrafish development: It’s in the maternalgenes. Curr. Opin. Genet. Dev. 19, 396–403. doi: 10.1016/j.gde.2009.06.002.

Arasu, A., Kumaseran, V., Sathyamoorthi, A., Palanisamy, R., Prabha, N., Bhatt, P., Roy, A., Thirumalai, M.K., Guanam, A. J., Pasupuleti, M. et al. (2013). Fish lily type lectin-1 contains β-prism 2 architecture: immunological characterization. Mol. Immunol. 56, 497–506.

Beishline, K. and Azizkhan-Clifford, J. (2015). Sp 1 and the ‘hallmarks of cancer’. FEBS J. 282, 224–258.

Bergsson, G., Agerberth, B., Jörnvall, H. and Gudmundsson, G.H. (2005). Isolation and identification of antimicrobial components from the epidermal mucus of Atlantic cod (*Gadus morhua*). FEBS J. 272, 4960–4969.

Bildfell, R. J., Markham, R. J. and Johnson, G. R. (1992). Purification and characterization of a rainbow trout egg lectin. J. Aquat. Anim. Health 4, 97–105.

Birkemo, G. A., Luders, T., Andersen, O., Nes, I. F. and Nissen-Meyer, J. (2003). Hipposin, a histone-derived antimicrobial peptide in Atlantic halibut (*Hippoglossus hippoglossus* L.). Biochim. Biophys. Acta 1646, 207–215.

Bouvet, J. (1976). Enveloping layer and the periderm of the trout embryo (*Salma Trutta fario* L.). Cell Tissue Res. 170, 367–380.

Cerliani, J. P., Stowell, S. R., Mascanfroni, I. D., Arthur, C. M., Cummings, R. D. and Rabinovich, G. A. (2011). Expanding the universe of cytokines and pattern recognition receptors: galectins and glycans in innate immunity. J. Clin. Immunol. 31, 10–21. doi: 10.1007/s10875-010-9494-2.

Chang, W-J. and Hwang, P-P. (2011). Development of Zebrafish Epidermis. Birth Defects Research (part C) 93, 205–214.

Chen, C. K., Chang, N. L. and Wang, A. H. (2011). The many blades of the beta-propeller proteins: conserved but versatile. Trends Biochem. Sci. 36, 553–561.

Chen, D., Fan, W., Lu Y., Ding, X., Chen, S. and Zhong, Q. (2012). A mammalian autophagosome maturation mechanism mediated by TECPR1 and the Atg12-Atg5 conjugate. Mol. Cell. 9, 629–41. doi: 10.1016/j.molcel.2011.12.036. Epub 2012 Feb 16.

Cordero, H., Brinchmann, M. F., Cuesta, A., Meseguer, J, and Esteban, M. A. (2015). Skin mucus proteome map of European sea bass (*Ditentrarchas labrax*). Proteomics 15, 4007–4020.

Cummings, R.D. (2014). If it is methylated it must be Tectonic. Proc. Natl. Acad. Sci. USA 27, 9669–9670. doi/10.1073/pnas.1408652111

Dash, S., Das, S. K., Samal, J. and Thatoi, H. N. (2018). Epidermal mucus, a major determinant in fish health: a review. Iran. J. Vet. Res. 19, 72–81.

De la Paz, J. F., Anguita, C., Diaz-Celis, C., Chavez., F. P., and Allende, M. L. (2020). Biomolecules 10, 1274. doi:10.3390/biom10091274

DiMichele, L. and Powers, D.A. (1984). Developmental and oxygen consumption rate differences between lactate dehydrogenase-B genotypes of *Fundulus heteroclitus* and their effect on hatching time. Physiol. Zool. 57, 52–56.

Dong, C. H., Yang, S. T., Yang, Z. A., Zhang, L. and Cui, J. F. (2004). A C-type Lectin associated and translocated with cortical granules during oocyte maturation and fertilization in fish. Dev. Biol. 265, 341–354.

Dorrington, M. G. and Fraser, I. D. C. (2019). NF-κB Signalling in Macrophages: Dynamics, Crosstalk, and Signal Integration. Frontiers in Immunol. 10, 705. doi: 10.3389/fimmu.2019.00706.

Elliot, D.G. (2011). Functional Morphology of the Integumentary System in Fishes. In Encyclopedia of Fish Physiology (ed. A. P. Farrell), pp. 476–488. New York. Academic Press.

Ellis, A. E. (2001). Innate Host defense mechanisms of fish against viruses and bacteria. Dev. Comp. Immunol. 25, 827–839.

Esteban, M. A. (2012). An overview of the immunological defenses in fish skin. ISRN Immunol. 2012, 1–30. Article ID 85470.3

Fast, M.D., Sims, D.E., Burka, J. F., Mustafa, A. and Ross, N. W. (2002). Skin morphology and non-specific defence parameters of mucus and plasma in the Rainbow trout (*Oncorhyncus mykiss*), coho and Atlantic salmon. Comp. Biochem. Phys. B. 132, 645–647.

Fraiberg, M., Tami-Yecheskel, B-C., Kokabi, K., Subic, N., Heimer, G., Eck, F., Nalbach, K., Behrends, C., Ben-Zeev, B. Shatz, O. and Elazar, S. (2021). Lysosomal targeting of autophagosomes by the TECPR domain of TECP2. Autophagy 17, 3096–3108.

Friedman, A. D. (2007). Transcriptional control of granulocyte and monocyte development. Oncogene 26, 6816–6828.

Fuentes, R. and Fernándes, J. (2010). Ooplasmic segregation in the zebrafish zygote and early embryo. Pattern of ooplasmic movement and transport pathways. Dev. Dyn. 239, 2172–2189. doi: 10.1002/dvdy.22349.

Ginzburg, A. S. (1968). Fertilization in Fishes and the problem of polyspermy. Israel Program for Scientific Translations. 1972, ISBN 7065 12391.

Groot, E.P. and Alderice, D.E. (1985). Fine structure of the external egg membrane of five species of Pacific Salmon and Steelhead trout. Can. J. Zool. 63, 552–566.

Guraya, S. S. (1986). The Cell and Molecular Biology of Fish Oogenesis. Monographs in Developmental Biology 18, pp. 1–223. New York. Karger.

Hamazaki, T.S., Iuchi, I and Yamagami, K. (1987). Isolation and partial characterization of a “spawning female-specific substance” in the teleost, *Oryzias latipes*. J. Exp. Zool. 242, 343–349.

Hamazaki, T.S., Nagahama, Y., Iuchi, I. and Yamagami, K. (1989). A glycoprotein from the liver constitutes the inner layer of the egg envelope (zona pellucida interna) of the fish, *Oryzias latipes*. Dev. Biol. 133, 101–110.

Hancock, R. E. W: and Scott, M. G. (2000). The role of antimicrobial peptides in animal defense. Proc. Natl. Acad. Sci. US. 97, 8856–8861.

Harstad, H., Lukacs, M.F., Bakke, H.G. and Grimholt, U. (2008). Multiple expressed MHC class II loci in salmonids; details of one non-classical region in Atlantic salmon (*Salmo salar*). BMC Genomics 28, 193. doi: 10.1186/1471-2164-9-193. PMID: 18439319; PMCID: PMC2386828.

Helvik, J.V. and Walther, B.T. (1992). Photo-regulation of the hatching process of halibut (*Hippoglossus hippoglossus*) eggs. J. Exp Zool. 263, 204–209.

Helvik, J.V. and Walther, B.T. (1993a). Development of hatchability in halibut (*Hippoglossus hippoglossus*) embryos. Int. J. Dev. Biol. 37, 487–490.

Helvik, J.V. and Walther, B.T. (1993b). Environmental parameters affecting induction of hatching in halibut (*Hippoglossus hippoglossus*) embryos. Mar. Biol. 116, 39–45.

Howe, K., Clark, M. D., Torroja, C. F., Torrance, J., Berthelot, C., Muffato, M., Collins, J.E., Humphray, S., McLaren, K., Matthews, L. et al. (2013). The zebrafish reference genome sequence and its relationship to the human genome. Nature 496, 498–503. doi: 10.1038/nature12111. Epub 2013 Apr 17.

Huh, C. G., Aldrich, J., Mottahedeh, J., Kwon, H., Johnson, C., and Marsh, R. (1998). Cloning and characterization of Physarum polycephalum tectonins. Homologues of Limulus lectin L-6. J. Biol. Chem. 273, 6565–6574. doi: 10.1074/jbc.273.11.6565.

Hyllner, S.J., Oppen-Berntsen, D.O., Helvik, J.V., Walther, B.T. and Haux, C. (1991). Estradiol-17β induces the major vitelline envelope proteins in both sexes in teleosts. J. Endocrinol. 131, 229–236.

Ingram, G. A. (1980). Substances involved in the natural resistance of fish to infection. A review. J. Fish Biol. 16, 23–60.

Inohaya, K., Yasumasu, S., Araki, K., Naruse, K., Yamazaki, K., Yasumasu, I., Iuchi, I. and Yamagami, K. (1997). Species-dependent migration of fish hatching gland cells that express astacin-like proteases. Develop. Growth Differ. 39, 191–197.

Jung, W. K., Kim, S. K. and Park, P. J. (2003). Purification and Characterization of a new lectin from the hard roe of skipjack tuna, *Katsuwonus pelamus*. Int. J. Biochem. Cell Biol. 35, 255–265.

Kane, D. A and Kimmel, C. B. (1993). The zebrafish mid-blastula transition Development 119, 447–456.

Kilpatrick, D.C. (2002). Animal Lectins: a historical introduction and overview. Biochim. Biophys. Acta 1572, 187–197.

Kunz, Y. (2004). *Developmental Biology of Teleostean Fishes*. Dordrecht, The Netherlands. Springer Verlag. ISBN 1-4020-2996-9.

Lien, S., Koop, B. F., Sandve, S. R., Miller, J.R., Kent, M. P., Nome, T., Hvidsten, T.R., Leong, J. S., Minkley, D. R., Zimin, A. et al. (2016). The Atlantic salmon genome provides insights into rediploidization. Nature 533, 200–205. https://doi.org/10.1038/nature17164

Lønning, S., Kjørsvik, E. and Davenport, J. (1984). The hardening process of the egg chorion of the cod. J. Fish Biol. 24, 505–522.

Low, D. H. P., Frecer, V., Le Saux, A., Srinivasan, G.A., Ho, B., Chen, J. and Ding, J.L. (2010). Molecular interfaces of the galactose-binding protein Tectonin domains in host-pathogen interaction. J. Biol. Chem. 285, 9898–9907. doi: 10.1074/jbc.M109.059774

Luders, T., Birkemo, G. A., Nissen-Meyer, J., Andersen, O. and Nes, I. F. (2005). Proline-conformation dependent antimicrobial activity of a ppline-rich histone H1 N-terminal peptide isolated from the skin mucus of Atlantic salmon. Antimicrob. Agents 49, 2399–2406.

Lund, V. and Olafsen, J. A. (1998). A comparative study of pentraxin-like proteins in different fish species. Dev. Comp. Immunol. 22, 185–194.

Magnadottir, B., Lange, S., Gudmundsdottir, S., Bøgwald, J. and Dalmo, B. A. (2005). Ontogeny of humoral immune parameters in fish. Fish Shellfish Immunol. 19, 429–439.

Matsushita, M., Matsushita, A., Endo, Y., Nakata, M., Kojima, N., Mizuochi, T. and Fujita, T. (2004). Origin of the classical complement pathway: lamprey orthologue of mammalian C1q acts as a lectin. Proc. Natl. Acad. Sci. US. 101, 10127–10131.

Mei, W., Lee, K. W., Marlow, F. L., Miller, A. L. and Mullins, M. C. (2009). hnRNP I is required to generated the Ca^2+^ signal that causes egg activation in zebrafish. Development 136, 2007–2017. doi: 10.1242/dev.037879.

Miftari, M.H. (2009). Leukolectins: Novel and Evolutionarily Conserved Proteins with Relevance for Vertebrate Macrophages. PhD-thesis, University of Bergen, Norway. ISBN 978-82-308-0842-9.

Miftari, M.H. and Walther, B.T. (2009). Leukolectin is a novel, unique and conserved protein in the PRR family. World Immune Regulation Meeting WIRM IV, Abstract March 29, Adaptive immunoregulatory mechanisms, Switzerland.

Modig, C., Westerlund, L. and Olsson, P. E. (2007). Oocyte zona pellucida proteins. In The Fish Oocyte (ed. Babin, P.J., Cerdà, J., Lubzens, E.) pp. 111–139. Dordrecht, The Netherlands. Springer Verlag. https://doi.org/10.1007/978-1-4020-6235-3_5

Oppen-Berntsen, D. O. (1990a). Oogenesis and Hatching in Teleostean Fishes, with special reference to Eggshell Proteins. *PhD Thesis*, University of Bergen, Norway.

Oppen-Berntsen, D.O., Helvik, J.V. and Walther, B.T. (1990b). The major structural proteins of cod (*Gadus morhua*) eggshells and protein crosslinking during teleost egg hardening. Dev. Biol. 137, 258–265.

Oppen-Berntsen, D.O., Bogsnes, A. and Walther, B.T. (1990c). The effects of hypoxia, alkalinity and neurochemicals on hatching of Atlantic Salmon (*Salmo salar*) eggs. Aquaculture 86, 417–430.

Oppen-Berntsen, D.O., Gram-Jensen, E. and Walther, B.T. (1992). Zona radiata proteins are synthesized by rainbow trout (*Oncorhyncus mykiss*) in response to oestradiol-17β. J. Endocrinol. 135, 293–302.

Oz-Levi, D., Ben-Zeev, B., Ruzzo, E. K., Hitomi, Y., Gelman, S., Pelak, K., Anikster, Y., Reznik-Wolf, H., Bar-Joseph, I., Olender, T., et al. (2012). Mutations in TECPR2 reveals a role for autophagy in hereditary spastic paraparesis. Am. J. Human Genetics 91, 1065–1072.

Pasquier, J., Cabau, C., Nguyen, T., Jouanno, E., Severac, D., Braasch, I., Journot, L., Pontarotti, P., Klopp, C., Postlethwait, J.H., et al. (2016). Gene evolution and gene expression after whole genome duplication in fish: the PhyloFish database. BMC Genomics 17, 368. doi: 10.1186/s12864-016-2709-z.

Pelka, K.E., Henn, K., Keck, A., Sapel, B. and Braunbeck, T. (2017). Size does matter-determination of the critical molecular size for the uptake of chemicals across the chorion of the zebrafish (*Danio rerio*) embryos. Aquat. Toxicol. 185, 1–10.

Qiao, H., Wang, Y., Zhang, X., Lu, R., Niu, J., Nan, F., Ke, D., Zeng, Z., Wang, Y. and Wang, B. (2022). Cross-species opsonic activity of zebrafish fish-egg lectin on mouse macrophages. Dev Comp Immunol. 129, 104332. doi: 10.1016/j.dci.2021.104332. Epub 2021 Dec 12. PMID: 34910945.

Rajan, B. J. M., Fernandes, C. M., Caipang, V., Kiron, J. H., Rombout, J. H. and Brinchmann, M. N. (2011). Proteome reference map of the skin mucus of Atlantic cod (*Gadus morhua*) revealing immune competent molecules. Fish Shellfish Immunol. 31, 224–231.

Rakers, S., Niklasson, L., Steinhagen, D., Kruse, C., Schauber, J., Sundell, K. and Paus, R. (2013). Antimicrobial peptides (AMPs) from fish epidermis: perspectives for investigative dermatology. J. Invest. Dermatol. 133, 1140–1149.

Rao, V., Marimuthu, K., Kupusamy, T., Rathinam, X., Arasu, M. V., Al-Dhabi, N. A. and Arochkiara, J. (2015). Defense properties in the epidermal mucus of different freshwater fish species. AACL Bioflux 8, 184–195.

Rong, C.J. (1997). Zonase from *Salmo salar*. Characterization of Endoproteases that Mediate Hatching Transition in the Sexual Life Cycle. *PhD Thesis*, University of Bergen, Norway. ISBN: 82–7653–010-9

Roussel, P. and Delmotte, P. (2004). The diversity of epithelial secreted mucins. Curr. Org. Chem. 8, 413–437.

Schoots, A. F. M., Evertse, P. A. C. M. and Denucé, J. M. (1983a). Ultrastructural Changes in hatching-gland cells of pike embryos (*Esox lucius* L.) and evidence for their degeneration by apoptosis. Cell Tissue Res. 229, 573–589.

Schoots, A.F.M., Meijer, R.C. and Denucé, J. M. (1983b). Dopaminergic regulation of hatching in fish embryos. Dev. Biol. 100, 59–63.

Sharon, N. and Lis, H. (2004) History of Lectins: From Hemagglutinins to Biological Recognition molecules. Glycobiology 14, 53R–62R. doi: 10.10931/glycob/cwh 122.

Sommer, R., Makshkova, O. N., Wohlschlager, T., Titz, A., Künzler, Verrot, A. (2018). Crystal structures of fungal Tectonin in Complex with O-Methylated Glycans suggest Key Role in Innate immune Defense. Structure 26, 391–402.

Stowell, S. R. C. M., Arthur, R., McBride, O., Berger, N., Razi, J., Heimburg-Molinaro, L. C., Rodrigues, J-P., Gourdine, A. J, Noll, S., von Gunten, D. F. et al. (2014). Microbial glycan microarrays define key features of host-microbial interactions. Nat. Chem. Biol. 10, 470–476. doi: 10.1038/nchembio.1525.

Subramanian, S., MacKinnon, S. l.. and Ross, N.W. (2007). A comparative study of innate immune parameters in the epidermal mucus of various fish species. Comp. Biochem. Phys. B 148, 356–263.

Tateno, H.I., Ogawa, T., Muramoto, K., Kamiya, H., Hirai, T. and Saneyoshi, M. (2001). A Novel Rhamnose-binding Lectin Family from Eggs of Steelhead Trout (*Oncorhynchus mykiss*) with Different Structures and Tissue Distribution. Bioscience, Biotechnology and Biochemistry 65, 1328–1338. doi.org/10.1271/bbb.65.1328

Tiralongo, F., Messina, G., Lombardo, B. M., Longhitano, L., Volti, G. L. and Tibullo, D. (2020). Skin Mucus of Marine Fish as a Source for the Develoment of Antimicrobial Agents. Frontiers in Marine Science 7, 541853. doi:10.3389/fmars.2020.541853.

Tsutsui, S. Tasumi, S. Suetake, H. and Suzuki, Y. (2003). Skin mucus lectin of pufferfish (*Fugu rubripes*) homologous to monocotyledonous plant lectin. J. Biol. Chem. 278, 20882–20889.

Walther, B.T. and Miftari, M.H. (2010) Leukolectins and Uses thereof. PCT patent WO2010049688, filed October 29, 2009, and published May 6, 2010.

Walther, B.T. and Rong, C.J. (1998). Non-Selfdegrading Endoprotease. PCT/NO98/00378. US Patent 6.346.245 (expired 2018).

Wang, Z. and Zhang, S. (2010) The role of lysozyme and complement in the antibacterial activity of zebrafish (Danio rerio) egg cytosol. Fish. Shellfish. Immun. 29, 773–777.

Wang, H., Ji, D., Shao, J. and Zhang, S. (2012). Maternal transfer and protective role of antibodies in zebrafish Danio rerio. Mol. Immunol. 51, 332–336.

Wang, Y., Bu, L., Yang, L., Li, H. and Zhang, S. (2016). Identification and functional characterization of fish-egg lectin in Zebrafish. Fish & Shellfish Immunol. dx.doi.org/j.fsi.2016.03.016

Wang, Z., Zhang, S., Tong, Z., Li, L. and Wang, G. (2020). Maternal Transfer and Protective Role of the Alternative Complement Components in Zebrafish Danio rerio. PLoS ONE. 4, e4498. Biomolecules 10, 1274 4498.

Wohlschlager, T., Butschi, A., Grassi, P., Sutov, G., Gaus, R., Hauck, D., Schmieder, S. S., Knobel, M., Titz, A., Dell, A. et al. (2014). Methylated glycans as conserved targets of animal and fungal innate defense. Proc. Natl. Acad. Sci. USA 111, E2787–2796.

Wu, M. H., Maier, E., Benz, R. and Hancock, R. E. W. (1999). Mechanism of interaction of different classes of cationic antimicrobial peptides with planar bilayers and with cytoplasmic membrane of *Escherichia coli*. Biochem. US. 38, 7235–7242.

Yamagami, K. (1988). Mechanisms of hatching in fish. In Fish Physiol. vol. XI, part A (ed. W. S. Hoar and D.J. Randall). pp. 447–499. New York: Academic Press.

Yokoya, S. and Ebina, Y. (1972). Hatching glands in salmonid fishes, *Salmo gairdneri*, *Salmo trutta, Salvelinus fontinalis* and *Salvelinus pluvius*. Cell Tissue Res. 172, 529–540.

